# Basal ganglia and cerebellar contributions to vocal emotion processing: a high resolution fMRI study

**DOI:** 10.1101/2020.11.26.399790

**Authors:** Leonardo Ceravolo, Sascha Frühholz, Jordan Pierce, Didier Grandjean, Julie Péron

## Abstract

Until recently, brain networks underlying emotional voice prosody decoding and processing were focused on modulations in primary and secondary auditory, ventral frontal and prefrontal cortices, and the amygdala. Growing interest for a specific role of the basal ganglia and cerebellum was recently brought into the spotlight. In the present study, we aimed at characterizing the role of such subcortical brain regions in vocal emotion processing, at the level of both brain activation and functional and effective connectivity, using high resolution functional magnetic resonance imaging. Variance explained by low-level acoustic parameters (fundamental frequency, voice energy) was also modelled. Wholebrain data revealed expected contributions of the temporal and frontal cortices, basal ganglia and cerebellum to vocal emotion processing, while functional connectivity analyses highlighted correlations between basal ganglia and cerebellum, especially for angry voices. Seed-to-seed and seed-to-voxel effective connectivity revealed direct connections within the basal ganglia ̶ especially between the putamen and external globus pallidus ̶ and between the subthalamic nucleus and the cerebellum. Our results speak in favour of crucial contributions of the basal ganglia, especially the putamen, external globus pallidus and subthalamic nucleus, and several cerebellar lobules and nuclei for an efficient decoding of and response to vocal emotions.

---

Social communication through voice entails semantic as well as prosodic meaning, the latter being generally defined as the melody of the human voice. The processing of human voice prosody leads to widespread changes in multiple cerebral regions, especially in the superior temporal and inferior frontal cortices (Ethofer, Anders et al. 2006, Frühholz and Grandjean 2013, Frühholz and Grandjean 2013, Grandjean in press). Although rarely put forward, the implication of the basal ganglia should be strongly emphasized. In fact, given their tripartite functional compartmentalization, whereby each basal ganglia (BG) is linked to either the motor, associative or limbic cortex (Alexander and Crutcher 1990, Lambert, Zrinzo et al. 2012), there is every reason to suppose that these structures play a major role in emotional processing in humans. This assertion is reinforced by both the BG’s intrinsic function and their functional and effective connectivity with the rest of the brain (Pierce and Péron 2020). There is growing evidence for the involvement of the BG in emotional processing, especially for emotions conveyed by the human voice (i.e., emotional prosody), not only directly, but also through their connections with structures known to be involved in emotional processing, such as the superior frontal and temporal gyri, the amygdala and the cerebellum (Thomasson, Saj et al. 2019). This involvement has been revealed by functional magnetic resonance imaging (fMRI) (Péron, Frühholz et al. 2016), electrophysiological data (Péron, Haegelen et al. 2014), lesion studies (Cohen, Riccio et al. 1994), as well as by deep brain stimulation of the BG, a neurosurgical technique that has recently drawn researchers’ attention to the possible functional roles of these structures in human emotional processing (for a review, see Péron, Frühholz et al. 2013).

Since the role of the BG was first hypothesized in vocal emotion decoding, evidence gathered from fMRI and lesion models has led to the hypothesis that they play a critical and potentially direct role in prosody processing, by promoting efficient decoding of emotional information from vocal cue sequences and rhythmic aspects of speech (Pell and Leonard 2003, Kotz and Schwartze 2010). The highly connected, closed loop nature of the BG make them perfectly situated to coordinate activity in other cortical and subcortical regions related to emotional voice perception. The BG, specifically the subthalamic nucleus (STN), may synchronize neural oscillations within a broader limbic network in order to facilitate efficient processing of auditory and emotion information (Péron, Frühholz et al. 2013). This synchronization would strengthen cortical representations of repeated stimulus-response pairings to form “chunks” of behavioural/cognitive response patterns that could be processed more automatically over learning (Graybiel 2008). Simultaneously, these chunks may be modified by the cerebellum to minimize the prediction error of an internal model based on its representation of the current sensory state and expected outcome of ongoing auditory processing (Sokolov, Miall et al. 2017, Bostan and Strick 2018). Furthermore, the BG and cerebellum may analyse temporal patterns in acoustic stimuli to extract salient emotional cues to feedback to cortex. Nevertheless, the way in which these subcortical and cortical structures exhibit coupling (or decoupling) in order to allow the emergence of a cognitive process such as emotional prosody recognition (i.e., functional integration) remains largely unexplored in affective neuroscience, especially the patterns of connectivity between the BG and the cerebellum, which should play a critical role in vocal emotion decoding (Thomasson, Saj et al. 2019, Pierce and Péron 2020). So far, we spoke about the BG as a whole concept without differentiating their abovementioned subparts and functional sub-territories. As for the subthalamic nucleus, the BG can be divided in at least three functional compartments relative to their cortical efferences: motor, associative and limbic (Alexander and Crutcher 1990, Lambert, Zrinzo et al. 2012, Pierce and Péron 2020). In the present study, we were specifically interested in the limbic BG due to the emotional nature of the stimuli presented to our participants. More specifically, BG regions of interest were the striatum, the globus pallidus (internal and external parts) and the subthalamic nucleus (Schneider, Habel et al. 2003, Wager, Barrett et al. 2008, Kotz, Schwartze et al. 2009, Péron, Frühholz et al. 2015, Pierce and Péron 2020). These BG regions also play a critical role in selecting a relevant response pattern nd inhibiting irrelevant ones̶ and in reward feedback and anticipation (Pierce and Péron 2020). BG efferences also connect them more directly to the cerebellum, which can also be separated into motor, associative, limbic and cognitive subparts (Leggio and Olivito 2018). Cerebellum functional subparts were recently highlighted by resting state functional connectivity (Buckner, Krienen et al. 2011), specific task-based parcellation (King, Hernandez-Castillo et al. 2019) and cerebellar topography (Leggio and Olivito 2018). In the scope of the present study, the cerebellum would help fine-tune the selected response initiated in the BG, generate an internal model of current goal states and somehow close the loop of reward encoding (Larry, Yarkoni et al. 2019, Pierce and Péron 2020) in addition to simultaneously assessing auditory timing for further iterations of vocal emotion decoding across time (Lesion studies: Grube, Cooper et al. 2010, Breska and Ivry 2016, Breska and Ivry 2018). Specific areas of the cerebellum associated with (vocal) emotion processing are the cerebellum crus of ansiform lobule I and II (Crus I,II), cerebellar lobules IV, V, VI, VIIb, VIII and IX, Vermis (Habas, Kamdar et al. 2009, Stoodley and Schmahmann 2009, Stoodley and Schmahmann 2009, Baumann and Mattingley 2012, Leggio and Olivito 2018, Thomasson, Saj et al. 2019, Pierce and Péron 2020) and deep cerebellar nuclei, especially the dentate (Pierce and Péron 2020) and fastigial nucleus (Wang, Dong et al. 2014, Zhang, Wang et al. 2016).

Although recent neuroimaging studies helped gain new insights into the role(s) of the BG in emotion processing, some of them still presented shortcomings that needed to be overcome. To date, these studies have failed to focus specifically on the BG, meaning that the measurement of the Blood-Oxygenation-Level Dependent (BOLD) signal was not restricted to these regions, thus reducing the spatial resolution in favour of a larger field of view. They also failed to investigate the functional and effective connectivity among the BG and between the BG and different subparts of the temporal regions (Frühholz, Ceravolo et al. 2011) that sustain emotional prosody processing, and more crucially between the BG and the cerebellum. Finally, the paradigms used so far in the literature did not test the impact of low-level acoustic parameters on voice prosody processing in the BG or cerebellum, even though these parameters greatly impact the BOLD signal at least in temporal and frontal brain regions (Schirmer and Kotz 2006, Frühholz, Ceravolo et al. 2012).

Considering abovementioned literature, the present study was designed to improve our current understanding of the functional integration of the BG and cerebellum during emotional prosody processing in humans, taking into account low-level acoustic parameters of interest such as synthesized fundamental frequency (*f*0) and energy, using high resolution fMRI in healthy participants. We therefore hypothesized: (i) an increase of BOLD signal in the STN, striatum, globus pallidus (internal, GPi; external, GPe) and cerebellum (Crus I-II, Vermis, cerebellar lobules IV-IX) during the processing of emotional (angry and happy) voices, as opposed to emotionally neutral voice prosody and (ii) similarly for emotional voices when removing variance explained by low-level acoustics (synthesized energy and *f0*); (iii) enhanced BOLD signal in the BG (STN, striatum, globus pallidus) for angry voice envelope (synthesized energy); (iiii) functional connectivity between the BG, especially in the STN and GPi/GPe, the cerebellum (Vermis and cerebellar lobules IV-IX, dentate nucleus) and temporal (superior temporal gyrus) and frontal voice areas (inferior frontal cortex, orbitofrontal cortex) when contrasting emotional to neutral voices (independently of synthesized energy and *f*0); (iiiii), enhanced effective coupling within the BG (striatum, STN, GPi/GPe) for angry and/or happy voices.

## Material and methods

### Participants

We initially included 19 healthy participants but excluded four of them from the analyses because of MRI signal artifacts (N=2) or psychiatric disorder (N=2). The remaining sample consisted of seven males and eight females (N=15), with a mean age of 30.5 years (SD *=* 3.48, range 27-37 years; mean age (SD) for female participants was 30.25 (3.24) and for male participants 30.85 (3.98)). All included participants were right-handed, native French speakers, and had normal or corrected-to-normal vision and normal hearing. None of them had a history of neurological disease or psychiatric disorder. Participants gave written informed consent for their participation in accordance with the ethical and data security guidelines of the University of Geneva. The study was approved by the local ethics committee and conducted according to the Declaration of Helsinki.

### Experimental setup

#### Main task

##### Stimuli

The vocal (prosodic) stimuli consisted of two pseudosentences spoken with different emotional prosodies (“*ne kali bam sud molen!*” and “*kun se mina lod belam?*”; mean duration = 1642 ms, range = 854-2788 ms) extracted from a previously validated database, the GEneva Multimodal Emotion Portrayals (GEMEP) corpus (Banziger and Scherer 2010). Alongside these prosodic stimuli (anger, happiness and neutral), we played synthesized stimuli, built from the original emotional and neutral sounds, in order to control for the temporal dynamics of energy and *f*0. These two basic acoustic features are known to be the most correlated with emotional prosody judgments (e.g., Banse and Scherer 1996, Grandjean, Banziger et al. 2006). The first type of synthetic stimulus (synthesized *intensity*) consisted of a section of white/pink noise, to which the intensity contour of the original stimulus was applied. The second type of synthetic stimulus (synthesized *f*0) was a series of pure sine waves (with constant amplitude), the frequency of which corresponded to the *f*0 of the original vocal stimulus, allowing us to maintain the temporal dynamics of the *f*0. Both synthetic stimuli had the same duration as in the original recordings. All sounds were matched for mean energy to avoid too strong loudness effects. Two runs were constructed, featuring the different kinds of stimuli in pseudorandom order (no more than three times for the same experimental condition). Each run contained 20 trials featuring anger stimuli, 20 trials featuring happiness stimuli, and 20 trials featuring neutral stimuli, as well as 15 synthesized intensity stimuli, 15 synthesized *f*0 stimuli, and one section of white noise at the beginning (first stimulus) with a gradual onset to accustom the participants to the auditory material. Each run contained a different list of stimuli. In each prosodic condition, we controlled for the pseudo-sentence being pronounced and the sex of the actor who pronounced the utterances: a female actor pronounced half the stimuli, half of them consisting of the pseudo-sentence “*ne kali bam sud molen!*”. The total duration of each run was ~10 minutes, and there was a short break between them. Each run contained pairs of identical subsequent stimuli, representing 10% of the total stimuli (pseudorandom order) to allow a one-back task to be performed by the participants, therefore forcing them to carefully attend each stimulus.

##### Experimental procedure, paradigm

In order to avoid expectancy effects, we varied in each trial the duration of the interval between the onset of the fixation cross and the onset of the auditory stimulus. In other words, the presentation of each auditory stimulus was preceded by a silent portion of pseudorandom duration, ranging from 50 to 250 ms, the so-called jitter (Fig.1). After the offset of the sound, we also included a silent portion ranging from 3000 to 3500 ms. In order to avoid the offset of the sound and the offset of the fixation cross being synchronous, we varied the duration of the interval between these two offsets. Finally, in order to minimize any retinal afterimage, we ensured that the color of the fixation cross did not contrast too greatly with the color of the desktop background.

For each trial, the participants were asked to keep their eyes open and relaxed. They were told they would hear meaningless speech uttered by male and female actors, as well as synthesized sounds. The binaurally recorded auditory stimuli were played through MR-compatible headphones. Loudness intensity was adjusted for each participant according to her/his hearing threshold at the beginning of the experiment. Participants were asked to focus on these auditory stimuli and to press a button whenever they heard two identical stimuli in a row. These one-back trials represented only 10% of all trials and were excluded from the analyses. The one-back task was administered to ensure that the patients were paying attention to the stimuli. Prior to the task, an MR-compatible response box (Current Designs Inc., Philadelphia, PA, USA) was placed beneath the participant’s fingers.

**Fig. 1:**
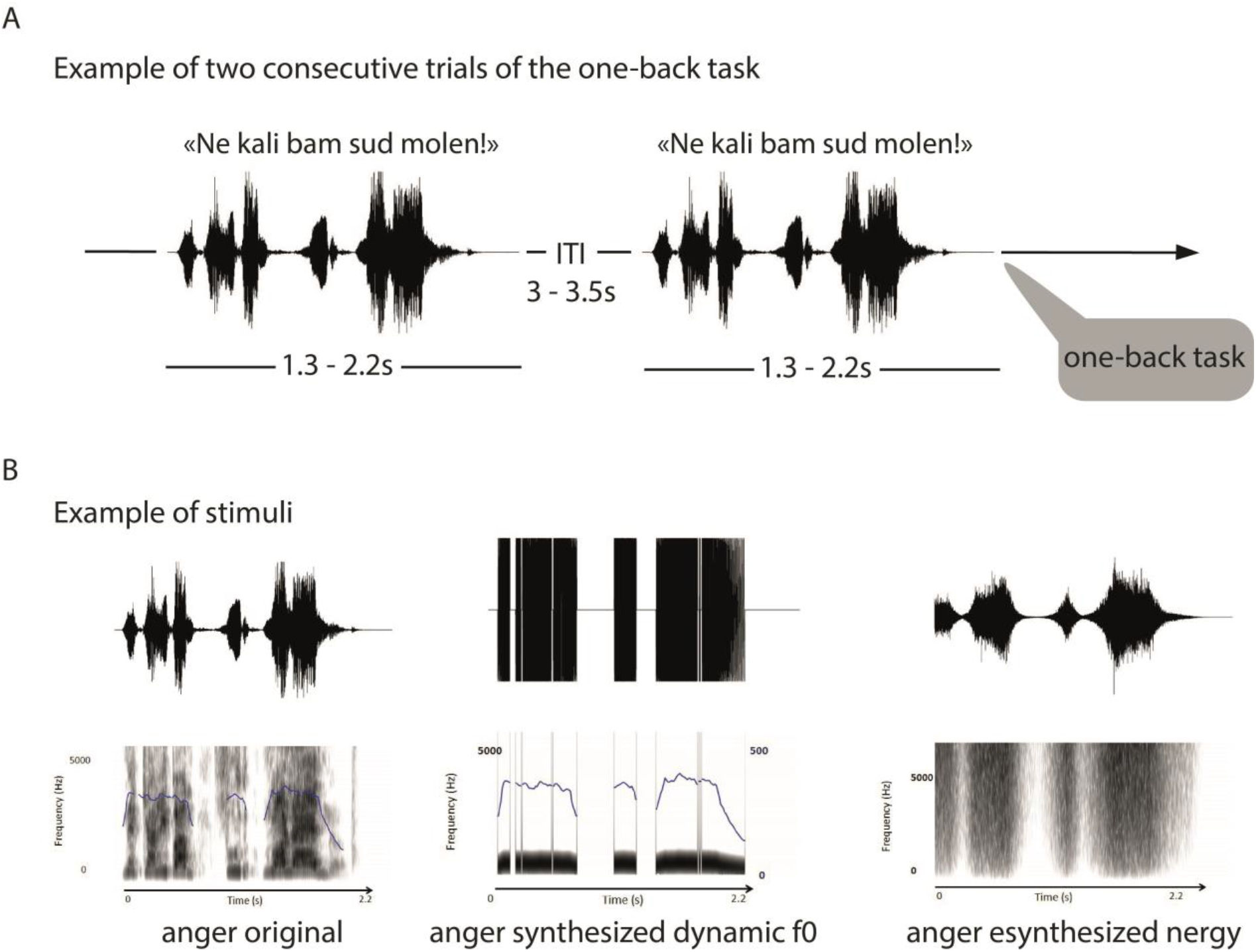
Experimental timeline and details of stimuli for the one-back task. A, Following technical scans (localizer and field map), the first run started for 10 min during which participants had to perform a one-back task on the voice presented auditorily to them using an MRI-compatible button box. The second run followed similarly for 10 more minutes and the session ended with the acquisition of an anatomical image for 5 min. During the complete session, the participant laid down in the scanner and had to pay attention to auditorily presented vocal stimuli and do a one-back task (10% of all trials). All stimuli had a duration of 1.3 to 2.2 s and an inter trial interval of 3 to 3.5 s. B, Voice stimuli consisted of pseudowords arranged in sentences with either original vocal signal, synthesized dynamic *f*0 manipulation or synthesized energy.

##### Image acquisition

Imaging was conducted at the Brain and Behaviour Laboratory (BBL) of the University of Geneva. For the main task, high-resolution imaging data was acquired on a 3T Siemens Trio System (Siemens, Erlangen, Germany) using a T2*-weighted gradient echo planar imaging sequence with 440 volumes per run (EPI; 1.5×1.5×2.2mm voxels, slice thickness=2mm, gap=0.2mm, 31 slices, RT=2320ms, TE=33ms, flip angle = 90°, matrix=128×128, field of view=192mm). The acquired volumes, representing a truncated field of view compared to standard wholebrain acquisition, were almost perpendicular to the anterior commissure-posterior commissure (AC/PC) line to cover all regions of interest, especially the basal ganglia, cerebellum and the temporal lobe (see Fig.S1 in the Supplementary material). Therefore, the term ‘wholebrain’ in this manuscript refers exclusively to our truncated field of view, not to volumes covering the wholebrain. The total number of volumes for our fifteen participants was 13’200 for a total number of slices of 409’200. A T1-weighted, magnetization-prepared, rapid-acquisition, gradient echo anatomical scan (slice thickness=1mm, 176 slices, RT=2530ms, TE=3.31ms, flip angle = 7°, matrix=256×256, FOV=256mm) was also acquired.

#### Image analysis

##### Wholebrain analyses

Functional images analysis was carried out using Statistical Parametric Mapping software 12 (SPM12, Wellcome Trust Centre for Neuroimaging, London, UK). Preprocessing steps included realignment to the first volume of the time series, slice timing, iterative normalization into the Montreal Neurological Institute space (Collins, Neelin et al. 1994) using the DARTEL toolbox (Ashburner 2007) and spatial smoothing with an isotropic Gaussian filter of 6 mm full width at half maximum. To remove low-frequency components, we used a high-pass filter with a cutoff frequency of 128 s. Anatomical locations were defined using a standardized coordinate database using the Automated Anatomical Labelling atlas (Tzourio-Mazoyer, Landeau et al. 2002) incorporated in the xjView toolbox (http://www.alivelearn.net/xjview), an atlas of the brainstem (Fonov, Evans et al. 2011), basal ganglia (Amunts, Lepage et al. 2013) and cerebellum (Diedrichsen, Balsters et al. 2009, Diedrichsen, Maderwald et al. 2011) displayed in FMRIB Software Library v6.0 (’FSL’; Smith, Jenkinson et al. 2004) through FSLeyes.

A general linear model was used to compute first-level statistics, in which each run was modelled as a distinct session and each trial was convolved with the hemodynamic response function, time-locked to the onset of each stimulus. Separate regressors were created for each condition, namely for the Emotion and the Acoustic Parameters factors (Design matrix columns for each run (N=9): anger original, anger *f*0, anger energy, happy original, happy *f*0, happy energy, neutral original, neutral *f*0, neutral energy). Finally, regressors of no-interest included the repetition trials of the one-back task that were concatenated across conditions and added as an additional regressor together with six motion parameters for each run to account for movement. Regressors of interest were used to compute nine simple contrasts (one per column of the design matrix, across runs) for each participant (across runs), leading to a main effect of each condition cited above at the first-level of analysis. Simple contrasts were then used in three distinct flexible factorial, second-level analyses. In model 1, the effect of the Emotion (angry, happy, neutral voices, acoustically untouched or ‘original’) factor was modelled with one Participant factor and one Emotion factor. In model 2, factors Participant, Emotion (angry, happy, neutral voices) and Acoustic Parameters (original, *f*0 synthesized, energy synthesized parameters) were included to model the two-way interaction between our main factors (Emotion*Acoustic Parameters). Model 3 included the main effect of the Acoustic Parameters (normal, *f*0 synthesized, energy synthesized parameters) factor, modelled with one Participant factor and one Acoustic Parameters factor. For each model, independence of the Participant factor was set to ‘true’, variance to ‘unequal’ and the Emotion, Acoustic Parameters and Emotion*Acoustic Parameters factors with independence as ‘false’, variance as ‘unequal’.

All neuroimaging activations were thresholded in SPM12 by using a wholebrain voxel-wise false discovery rate (FDR) correction at *p*<.05 with an arbitrary cluster extent of k>10 voxels.

##### Functional and effective connectivity analysis

Functional and effective connectivity analyses were performed using the CONN toolbox (Whitfield-Gabrieli and Nieto-Castanon 2012) version 18.b implemented in Matlab 9.0 (The MathWorks, Inc., Natick, MA, USA) for the two-way interaction between our factors, namely Emotion and Acoustic Parameters (design matrix identical to wholebrain analyses). As in wholebrain data analysis, repetition trials of the one-back task were modelled as a single column including a concatenation of all their onset times across conditions (regressor of no-interest). Functional connectivity analyses were mainly carried out to orient further effective connectivity analysis and we decided to report both types of connectivity for a clear overview of the results. Functional connectivity analyses were computed using as seeds each region of interest (ROI) of the following atlases: the Automated Anatomical Labelling atlas (’aal’; 58 ROI; Tzourio-Mazoyer, Landeau et al. 2002), an atlas of the brainstem (23 ROI; Fonov, Evans et al. 2011), basal ganglia (22 ROI; Amunts, Lepage et al. 2013) and cerebellum (34 ROI; Diedrichsen, Balsters et al. 2009, Diedrichsen, Maderwald et al. 2011). All ROI (N=137; Supplementary Table 1) were within the bounds of our truncated field of view. Frontal, parietal and occipital areas outside the bounds of our field of view, specifically of the ‘aal’ atlas, were isolated through CONN time-course visualization and removed from the analyses. For effective connectivity analyses and according to our hypotheses, seed regions were limited to the basal ganglia (22 ROI; Amunts, Lepage et al. 2013). Spurious sources of noise were estimated and removed using the automated toolbox preprocessing algorithm, and the residual BOLD time-series was band-pass filtered using a low frequency window (0.008 < f < 0.09 Hz). Correlation maps were then created for each condition of interest by taking the residual BOLD time-course for each condition from atlas regions of interest and computing bivariate Pearson’s correlation coefficients between the time courses of each voxel of each ROI of the atlas, averaged by ROI (‘functional connectivity’ analyses). ‘Effective connectivity’ was approached using multivariate regressions between each seed ROI and all other ROI ̶ or all brain voxels for seed to voxel analysis ̶ and a model was generated and used to characterize the direct connectivity between pairs. For both types of connectivity, we used generalized psychophysiological interaction (gPPI) measures, representing the level of task-modulated (often labelled ‘effective’) connectivity between ROI or between ROI and voxels. gPPI is computed using a separate multiple regression model for each target (ROI/voxel). Each model includes three predictors: 1) task effects convolved with a canonical hemodynamic response function (psychological factor); 2) each seed ROI BOLD time series (physiological factor) and 3) the interaction term between the psychological and the physiological factors, the output of which is regression coefficients associated with this interaction term. Finally, group-level analyses were performed on these regression coefficients to assess for main effects within-group for contrasts of interest in seed-to-seed and seed-to-voxel analyses. Therefore, ‘functional connectivity’ is defined in the present study as a gPPI analysis using bivariate correlations between ROI, while ‘effective connectivity’ defines the gPPI analysis using multivariate regressions between ROI/voxels. Connectivity analyses were computed using methods in line with most recent best practices (Reid, Headley et al. 2019). For both analyses, type I error was controlled by the use of seed-level (seed-to-seed analyses) and cluster-level (seed-to-voxel analysis) false discovery rate correction with *p*< .05 FDR to correct for multiple comparisons.

## Results

### Wholebrain results

We performed voxel-level general linear analyses subdivided into three different models in order to find enhanced brain activity related to the factorial design of our data. The models of interest were model 1 and 2, in which we modelled the Emotion factor and the two-way interaction between Emotion and Acoustic Parameters factors. The former analysis revealed emotion-specific enhanced patterns of activity that are presented in this section (for the general effect of Emotion, see Fig.S2 in the Supplementary material), while the full interaction between factors did not yield any significant results. We present, however, one significant result of interest, as part of our hypotheses, for the rhythmicity of angry voices (synthesized energy of angry > neutral prosody). Finally, results for model 3 – the main effect of Acoustic Parameters – are reported in the supplementary data (Supplementary data, Tables 2-4).

### Main effect of Emotion factor

Wholebrain results for the Emotion factor revealed significant enhanced activity for both angry > neutral voices (Table 1) and happy > neutral voices (Table 2) contrasts. Enhanced activations for emotional (angry and happy) compared to neutral voices were also significant especially in the superior temporal cortex and inferior frontal cortex, bilaterally (see Table 3). Brain activity specific to angry voices (angry > neutral voices) replicated the involvement of the temporal cortex for processing such stimuli, especially in the anterior part of the middle temporal cortex (aMTG) and the posterior superior temporal gyrus and sulcus (pSTG and pSTS, respectively), bilaterally (Fig.2ABG). Enhanced activity was also observed in medial brain areas such as the anterior cingulate cortex (ACC), the parahippocampal gyrus and the fusiform gyrus (Fig.2CD). Activity in the basal ganglia was restricted to the external globus pallidus (GPe) while we also observed enhanced activity in several parts of the thalamus (Fig.2E). Finally, large parts of the cerebellum were also more active (Fig.2G) during angry as opposed to neutral voice processing, namely the Crus II area (Fig.2B), lobules IV-V and VI (Fig.2CF), Vermis area VI (Fig.2D) as well as deep nuclei such as the dentate (Fig.2CF) and fastigial nucleus (Fig.2F). More details are available in Table 1.

**Table 1:** Activations, cluster size and coordinates for angry > neutral voices contrast, wholebrain voxel-wise *p*<.05 FDR correction, k>10.

| <u>MNI coordinates</u> |  |  |  |  |  |  |
| --- | --- | --- | --- | --- | --- | --- |
| Region label | Hemisphere | X | Y | Z | T value | Cluster size (voxels) |
| Precentral gyrus | R | 60 | 8 | 26 | 12.77 | 11526 |
| Postcentral gyrus | R | -58 | -14 | 18 | 10.91 |  |
| Precentral gyrus | R | 52 | 2 | 46 | 10.88 |  |
| STG, posterior | R | 54 | -34 | 12 | 9.75 | 84 |
| STS, posterior | L | -64 | -48 | 6 | 9.04 | 110 |
| ITG, posterior | L | -46 | -36 | -16 | 8.71 | 85 |
| Amygdala | L | -26 | -6 | -24 | 6.99 | 84 |
| Supp Motor Area | R | 8 | 6 | 58 | 6.36 | 47 |
| Mid Frontal gyrus | L | -28 | 26 | 42 | 6.07 | 211 |
| Mid Frontal gyrus | L | -40 | 36 | 22 | 5.70 | 309 |
| Sup Frontal gyrus | L | -12 | 36 | 48 | 5.58 | 112 |
| Cereb Nucl Fastigial | R | 6 | -56 | -28 | 5.54 | 133 |
| ACC | R | 8 | 16 | 28 | 5.49 | 117 |
| Globus Pallidus | R | 18 | -4 | 8 | 5.48 | 105 |
| Cereb Crus 2 | R | 10 | -84 | -36 | 5.29 | 11 |
| Globus Pallidus | L | -18 | 0 | -4 | 5.07 | 44 |
| Temporal pole | L | -58 | 10 | -8 | 5.02 | 31 |
| ACC | R | 8 | 30 | 18 | 4.92 | 45 |
| IFG triangularis | L | -50 | 28 | 10 | 4.72 | 14 |
| Cereb Lob 8 | L | -28 | -58 | -52 | 4.59 | 29 |
| Supramarginal gyrus | L | -66 | -22 | 28 | 4.59 | 13 |
| IFG opercularis | L | -58 | 16 | 10 | 4.33 | 24 |
| Sup Frontal gyrus | R | 14 | 48 | 34 | 4.26 | 32 |
| IFG triangularis | L | -42 | 20 | 8 | 4.17 | 36 |
| Cereb Crus 2 | L | -40 | -74 | -44 | 4.14 | 10 |
| Putamen | L | -14 | 10 | -10 | 4.09 | 27 |
| Cereb Vermis 1-2 | L | 0 | -42 | -24 | 3.97 | 20 |
| Cereb Lob 6 | L | -30 | -54 | -38 | 3.96 | 26 |
| Sup Frontal gyrus, medial | L | -8 | 48 | 36 | 3.91 | 33 |
| ACC | L | -12 | 26 | 28 | 3.75 | 12 |
| Cereb Crus 1 | L | -40 | -76 | -26 | 3.74 | 18 |
| STG, mid | L | -54 | -10 | -4 | 3.59 | 12 |
| MTG, anterior | L | -62 | -8 | -18 | 3.28 | 20 |

L: left; R: right; MNI: Montreal neurological institute; STG: superior temporal gyrus; STS: superior temporal sulcus; ITG: inferior temporal gyrus; Supp Motor Area: supplementary motor area; Mid: middle; Sup: superior; Cereb Nucl Fastigial: fastigial nucleus of the cerebellum; ACC: anterior cingulate cortex; Cereb Crus: cerebellum crus of ansiform lobule; IFG: inferior frontal gyrus; Cereb Lob: cerebellum lobule; Cereb: cerebellum; MTG: middle temporal gyrus.

**Table 2:**
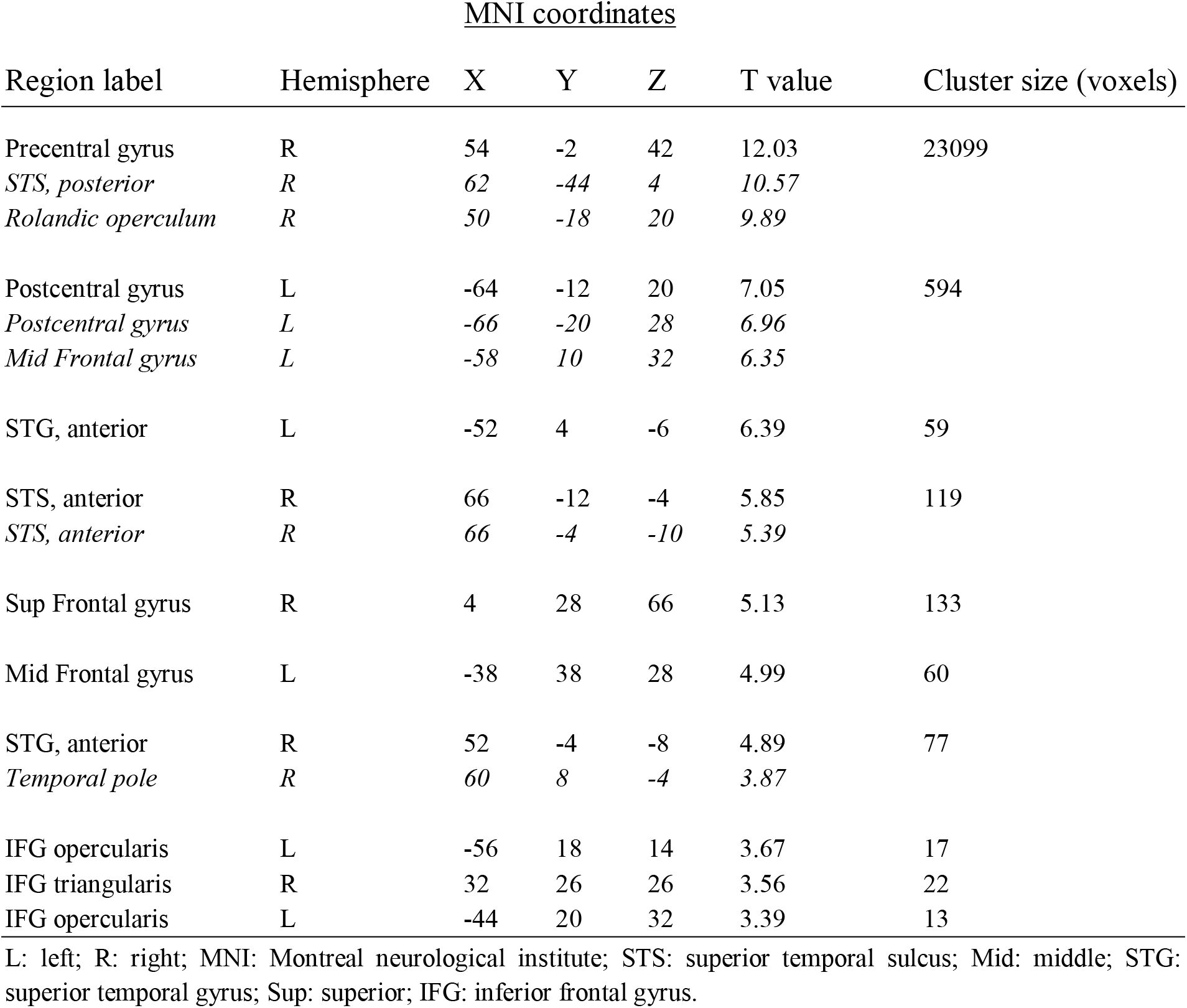
Activations, cluster size and coordinates for happy > neutral voices contrast, wholebrain voxel-wise *p*<.05 FDR correction, k>10.

**Table 3:**
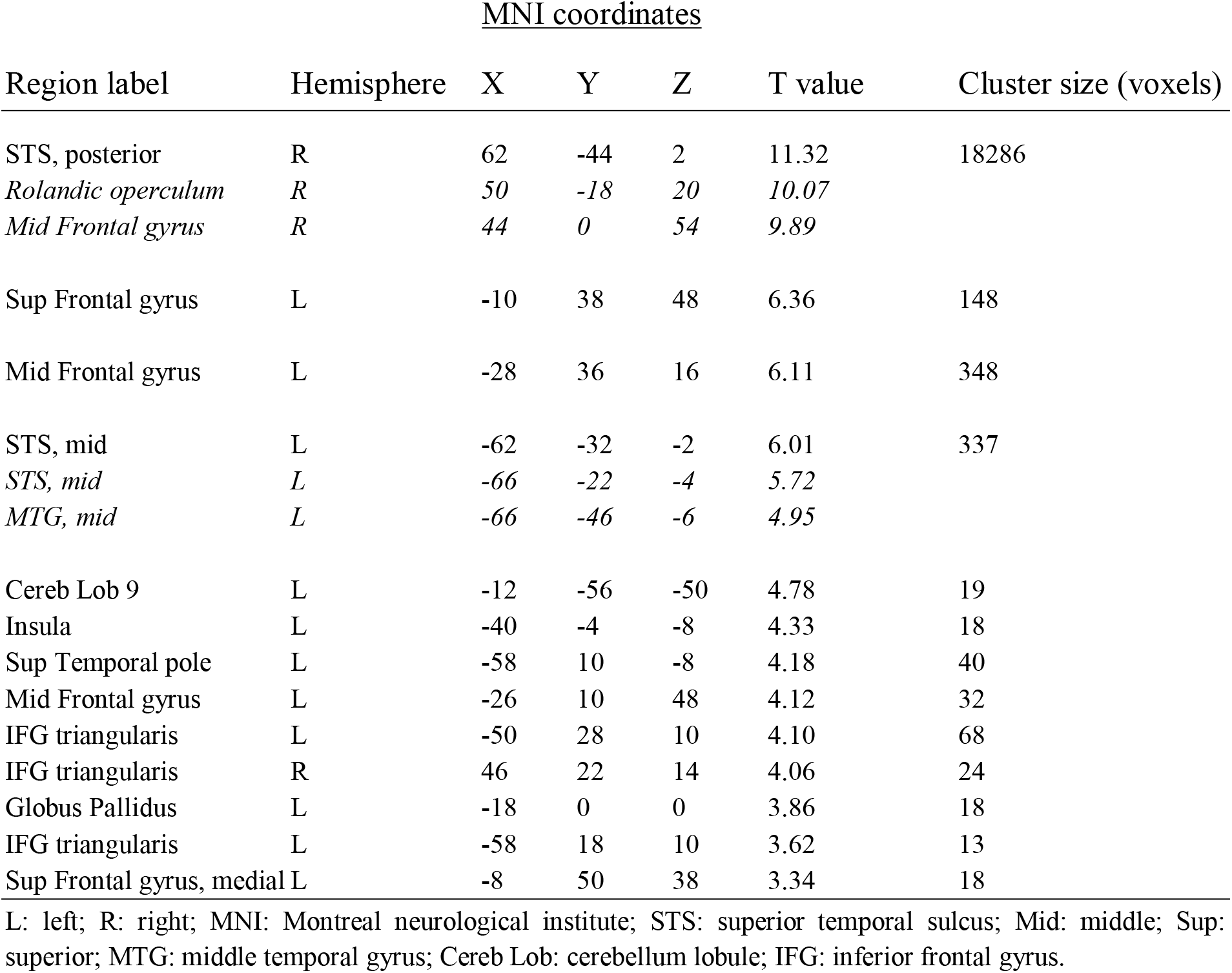
Activations, cluster size and coordinates for angry & happy> neutral voices contrast, wholebrain voxel-wise *p*<.05 FDR correction, k>10.

**Fig. 2:**
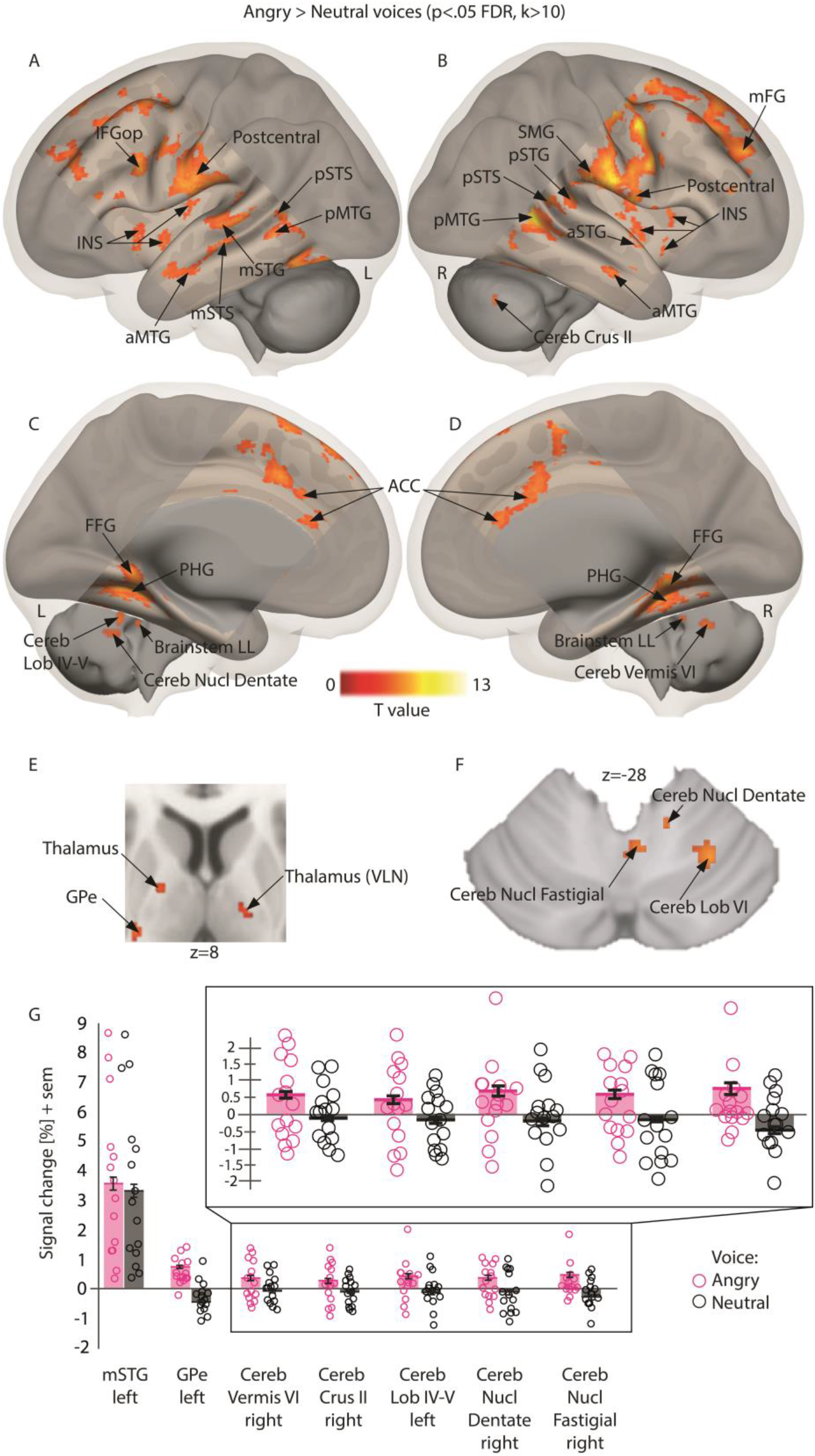
Enhanced brain measures for implicitly processing angry compared to neutral voices, corrected for multiple comparisons (wholebrain voxel-wise *p*<.05 FDR, k>10 voxels). A-B, Lateral activations rendered on a sagittal image highlighting middle and superior temporal regions. C-D, Medial activations of the anterior cingulate cortex, parahippocampal cortex and cerebellum. E, Subcortical activity in the thalamus and globus pallidus displayed on an axial slice. F, Cerebellar activations displayed on an axial slice. G, Percentage of signal change extracted using singular value decomposition on 9 voxels around each peak in a subset of regions with individual values (circles), mean values (bars) and standard error of the mean (error bars) for angry and neutral voices. Pink circles: angry voices; Black circles: neutral voices. L: left; R: right; IFGop: inferior frontal gyrus pars opercularis; STG: superior temporal gyrus; STS: superior temporal sulcus; MTG: middle temporal gyrus; INS: insula; SMG: supramarginal gyrus; FG: frontal gyrus; FFG: fusiform gyrus; PHG: parahippocampal gyrus; ACC: anterior cingulate cortex; Cereb: cerebellum; Cereb Lob: cerebellum lobule; Cereb Nucl Dentate: dentate nucleus of the cerebellum; Cereb Nucl Fastigial: fastigial nucleus of the cerebellum; Brainstem LL: lateral lemniscus of the brainstem; Thalamus VLN: ventral lateral nucleus of the thalamus; GPe: external globus pallidus; Cereb Crus: cerebellum crus of ansiform lobule; ACC: anterior cingulate cortex. ‘a’ prefix: anterior part; ‘m’ prefix: mid part; ‘p’ prefix: posterior part.

As for angry voices, brain activity specific to normal happy voices (happy > neutral voices) highlighted the anterior, mid and posterior portions of the temporal cortex (aSTS, aMTG; mSTS, mSTG; pSTS, pSTG, pMTG, respectively), bilaterally (Fig.3ABG). Enhanced activity was medially observed in the ACC, parahippocampal gyrus and fusiform gyrus (Fig.3CD). Increase of activity in the basal ganglia was observed in the GPe and bilateral putamen, and in the ventral lateral nucleus of the thalamus (Fig.3E). Multiple subparts of the cerebellum showed significant differences. Cerebellum areas were more activated (Fig.3G) during happy as opposed to neutral voice processing, especially in the lateral Crus I area, bilaterally (Fig.3ABF), in lobules VI, VIIb and VIII (Fig.3CDF), in Vermis areas III and IV-V (Fig.3CD) as well as in the dentate nucleus (Fig.3F). More details are available in Table 2.

**Fig. 3:**
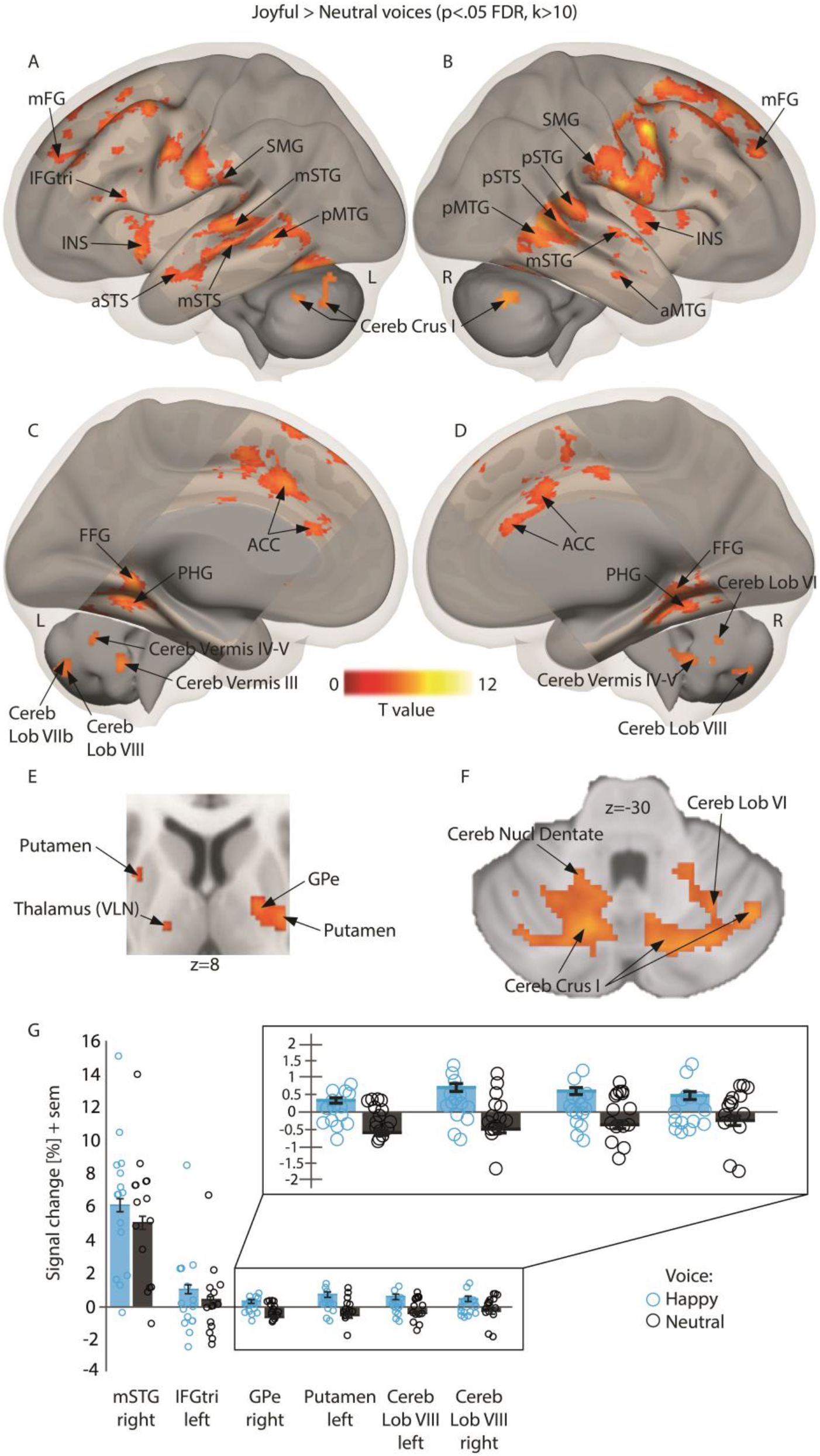
Enhanced brain measures for implicitly processing happy compared to neutral voices, corrected for multiple comparisons (wholebrain voxel-wise *p*<.05 FDR, k>10 voxels). A-B, Lateral activations rendered on a sagittal image highlighting middle, superior temporal and cerebellar regions. C-D, Medial activations of the anterior cingulate cortex, parahippocampal cortex and cerebellum. E, Subcortical activity in the putamen, thalamus and globus pallidus displayed on an axial slice. F, Cerebellar activations displayed on an axial slice. G, Percentage of signal change extracted using singular value decomposition on 9 voxels around each peak in a subset of regions with individual values (circles), mean values (bars) and standard error of the mean (error bars) for happy and neutral voices. Blue circles: happy voices; Black circles: neutral voices. L: left; R: right; IFGtri: inferior frontal gyrus triangularis part; STG: superior temporal gyrus; STS: superior temporal sulcus; MTG: middle temporal gyrus; INS: insula; SMG: supramarginal gyrus; FG: frontal gyrus; FFG: fusiform gyrus; PHG: parahippocampal gyrus; ACC: anterior cingulate cortex; Cereb: cerebellum; Cereb Lob: cerebellum lobule; Cereb Nucl Dentate: dentate nucleus of the cerebellum; Brainstem LL: lateral lemniscus of the brainstem; Thalamus VLN: ventral lateral nucleus of the thalamus; GPe: external globus pallidus; Cereb Crus: cerebellum crus of ansiform lobule; ACC: anterior cingulate cortex. ‘a’ prefix: anterior part; ‘m’ prefix: mid part; ‘p’ prefix: posterior part.

### Interaction effect between Emotion and Acoustic Parameters factors

The full, two-way interaction between our Emotion and Acoustic Parameters factors did not reveal significant results when contrasting angry or happy voices to neutral voices while taking into account normal compared to synthesized voices. We, however, had a specific hypothesis concerning the rhythmicity of angry voices, namely the impact of the ‘envelope’ of such voices on basal ganglia regions. We therefore used model 3 to compute a contrast dedicated to highlighting brain regions sensitive to the envelope of angry compared to neutral, synthesized energy voices [synthesized energy for angry > neutral voices]. The contrast revealed enhanced activity in the left ventral lateral and lateral posterior nucleus of the thalamus, putamen, substantia nigra, right caudate head, thalamus as well as in the bilateral insula, left amygdala and right mid-to-anterior and posterior STG (Table 4). Similar regions, especially large parts of the STG and STS, were also more active for the synthesized energy of happy voices, namely for the [synthesized energy for happy > neutral voices] contrast (Table 5).

**Table 4:** Activations, cluster size and coordinates for angry > neutral synthesized energy voices contrast, wholebrain voxel-wise *p*<.05 FDR correction, k>10.

| <u>MNI coordinates</u> |  |  |  |  |  |  |
| --- | --- | --- | --- | --- | --- | --- |
| Region label | Hemisphere | X | Y | Z | T value | Cluster size (voxels) |
| Precentral gyrus | R | 60 | 8 | 26 | 7.99 | 226 |
| Postcentral gyrus | L | -56 | -12 | 18 | 7.57 | 397 |
| MTG, posterior | R | 60 | -46 | 4 | 6.03 | 34 |
| Insula | R | 44 | -10 | 4 | 5.81 | 216 |
| STG, anterior | R | 56 | -6 | -4 | 5.23 |  |
| STG, posterior | R | 54 | -34 | 12 | 5.58 | 25 |
| STS, posterior | L | -62 | -48 | 6 | 5.43 | 20 |
| Hypothalamus | R | 2 | -4 | -10 | 5.20 | 53 |
| Amygdala | L | -26 | -6 | -24 | 5.12 | 28 |
| MTG, anterior | R | 66 | -4 | -16 | 4.89 | 24 |
| Parahippocampal gyrus | R | 26 | -38 | -16 | 4.72 | 151 |
| Substantia nigra | L | -8 | -28 | -6 | 4.71 | 12 |
| Parahippocampal gyrus | L | -24 | -34 | -4 | 4.04 | 17 |
| Thalamus, LPN | L | -20 | -16 | 10 | 3.65 | 10 |
| Caudate head | R | 4 | 10 | -6 | 3.43 | 15 |
| Putamen | L | -16 | 8 | -8 | 3.40 | 10 |

L: left; R: right; MNI: Montreal neurological institute; MTG: middle temporal gyrus; STG: superior temporal gyrus; STS: superior temporal sulcus; LPN: lateral posterior nucleus.

**Table 5:** Activations, cluster size and coordinates for happy > neutral synthesized energy voices contrast, *p*<.05 voxel-wise FDR correction, k>10.

| <u>MNI coordinates</u> |  |  |  |  |  |  |
| --- | --- | --- | --- | --- | --- | --- |
| Region label | Hemisphere | X | Y | Z | T value | Cluster size (voxels) |
| Precentral gyrus | R | 56 | -2 | 42 | 7.96 | 229 |
| STS, posterior | R | 62 | -44 | 4 | 6.82 | 486 |
| STS, posterior | L | -64 | -32 | -4 | 6.30 | 377 |
| Mid frontal gyrus | R | 32 | 28 | 48 | 5.76 | 663 |
| Putamen | R | 28 | -10 | 8 | 5.47 | 232 |
| Parahippocampal gyrus | L | -22 | -44 | -4 | 5.19 | 454 |
| IFG triangularis | L | -44 | 26 | 8 | 4.57 | 55 |
| ACC | L | -4 | 12 | 38 | 4.45 | 153 |
| Putamen | L | -28 | -18 | 10 | 4.41 | 33 |
| Amygdala | L | -26 | -8 | -22 | 4.20 | 63 |
| Thalamus, VPLN | L | -18 | -14 | 6 | 3.45 | 10 |
| Caudate head | L | -8 | 20 | 0 | 3.35 | 10 |

L: left; R: right; MNI: Montreal neurological institute; STS: superior temporal sulcus; Mid: middle; IFG: inferior frontal gyrus; ACC: anterior cingulate cortex; VPLN: ventral posterior lateral nucleus.

### Functional connectivity results

Wholebrain analyses revealed significant results for both of our factors (Emotion, Acoustic Parameters) but their interaction did not yield any above-statistical-threshold activations. Computing functional/effective connectivity analyses (both seed-to-seed and seed-to-voxel), however, did reveal several coupled and anti-coupled networks underlying such two-way interaction between the Emotion and the Acoustic Parameters factors. While functional connectivity results were primarily used to further compute effective connectivity, we kept them in the present section due to their specificity and general meaning. These results are presented below.

### Seed-to-seed functional connectivity

Computed using 137 ROI composed of 58 ‘aal’ regions within our field of view, 23 brainstem regions, 22 basal ganglia regions and 34 cerebellum regions, seed-to-seed analyses revealed significant results for the interaction between Emotion and Acoustic Parameters factors, for each emotion of interest. Our contrasts of interest therefore included angry or happy compared to neutral voices when spoken normally as opposed to synthesized *f*0 and energy voices. Seed-to-seed functional connectivity specific to angry original voices were therefore computed with the [angry > neutral voices * original > *f*0 & energy synthesized voices] contrast, revealing coupled networks. As predicted, we observed coupling between the basal ganglia and the cerebellum, more specifically between the left GPe and right cerebellum lobule X (Fig. 4). Coupled functional connectivity was also observed between the left pSTG and right frontal operculum and in the brainstem between major motor (right parieto-occipito-temporo-pontine tract) and sensory tracts (bilateral spinothalamic tract). Detailed results are reported in Table 6.

**Fig. 4:**
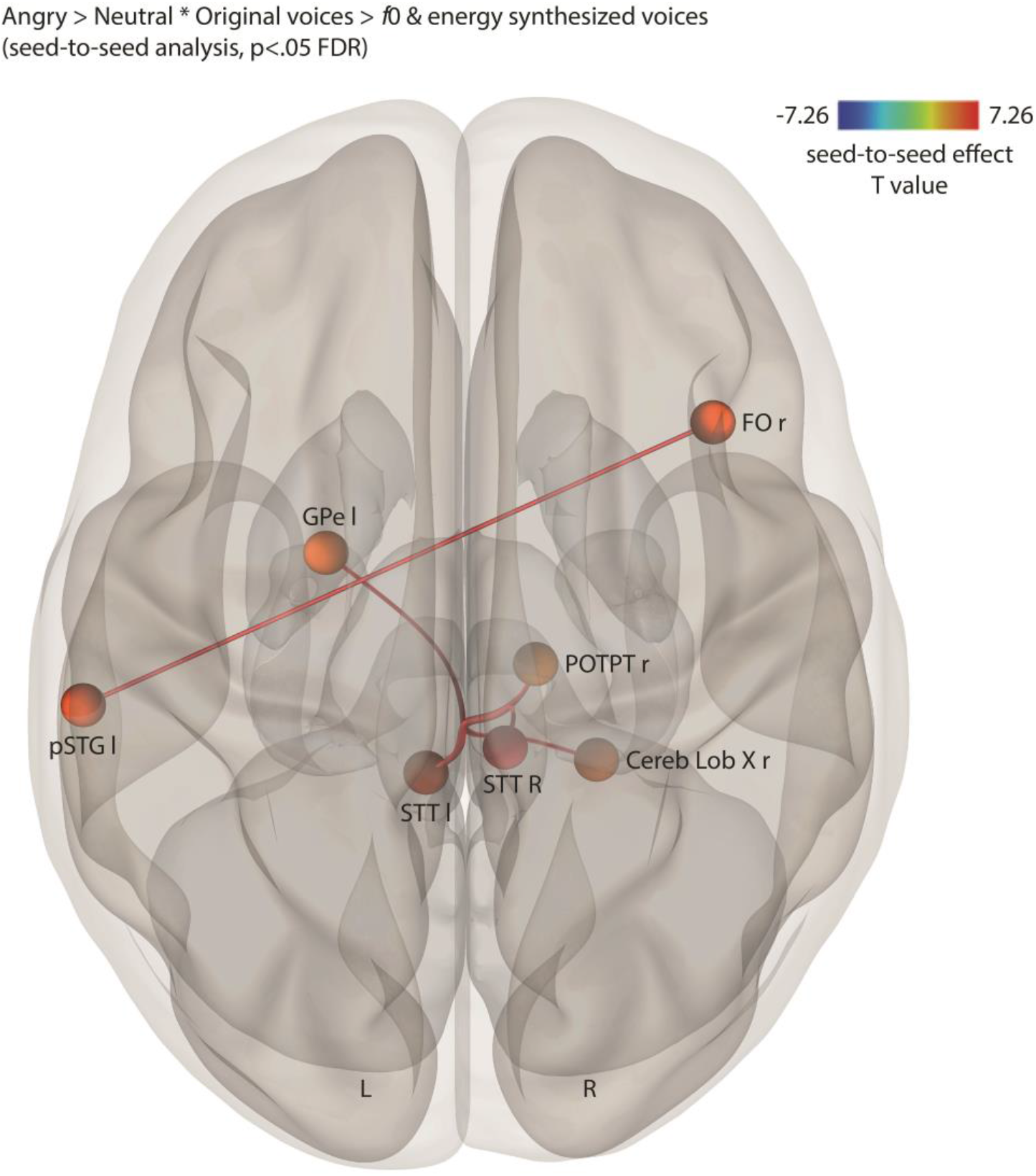
Coupled seed-to-seed, gPPI functional connectivity for the interaction between the Emotion and the Acoustic parameter factors contrasting angry > neutral voices * original > *f*0 & energy synthesized voices, corrected for multiple comparisons (*p*<.05 FDR). l and L: left; r and R: right; FO: frontal operculum; GPe: external globus pallidus; pSTG: posterior superior temporal gyrus; STT: spinothalamic tract of the brainstem; POTPT: parieto-occipito-temporo-pontine tract of the brainstem; Cereb Lob: cerebellum lobule.

**Table 6:** Seed-to-seed functional connectivity (gPPI) for angry > neutral normal > f0 & energy synthesized voices contrast, *p*<.05 seed-level FDR correction, two-tailed.

| 1018 | Seed region label | Target region label | T value | pFDR |
| --- | --- | --- | --- | --- |
| 1019 | pSTG l | FO r | 7.26 | 0.0006 |
| 1020 | GPe l | Cereb Lob X r | 4.69 | 0.0470 |
| 1021 | STT l | POTPT r | 5.71 | 0.0073 |
| 1022 | STT l | STT r | 5.30 | 0.0077 |

FO: frontal operculum; GPe: external globus pallidus; pSTG: posterior superior temporal gyrus; STT: spinothalamic tract of the brainstem; POTPT: parieto-occipito-temporo-pontine tract of the brainstem; Cereb Lob: cerebellum lobule; l: left; r: right.

Looking at positive emotion stimuli, happy voices yielded coupled and anti-coupled seed-to-seed functional connectivity results, as seen in the [happy > neutral voices * original > *f*0 & energy synthesized voices] contrast (Fig.5). Coupled functional connectivity revealed three distinct networks: 1) Internal globus pallidus (GPi) and aSTG in the right hemisphere; 2) Left pMTG and right central operculum cortex; 3) Right corticospinal tract (major motor tract) and right lateral lemniscus (major sensory tract). Happy voices also led to two separate anti-coupled networks involving the right paracingulate cortex and subcalcarine cortex as well as in posterior temporal areas, namely between the left pMTG and right pSTG (Fig.5). Details reported in Table 7.

**Fig. 5:**
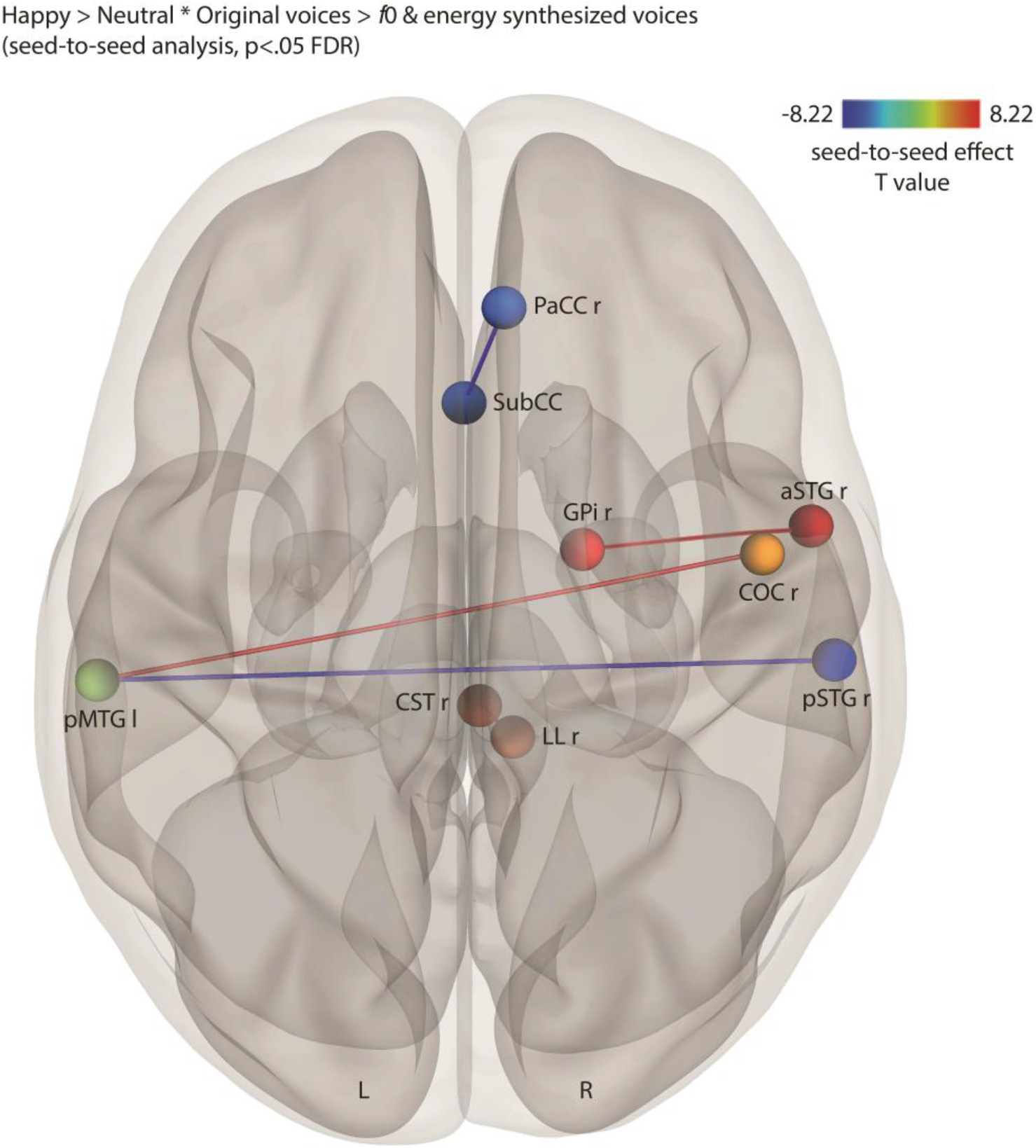
Coupled and anti-coupled seed-to-seed, gPPI functional connectivity for the interaction between the Emotion and the Acoustic parameter factors contrasting happy > neutral voices * original > *f*0 & energy synthesized voices, corrected for multiple comparisons (*p*<.05 FDR). l and L: left; r and R: right; PaCC: paracingulate cortex; SubCC: subcalcarine cortex; GPi: internal globus pallidus; COC: central operculum cortex; aSTG: anterior superior temporal gyrus; pSTG: posterior superior temporal gyrus; pMTG: posterior middle temporal gyrus; CST: corticospinal tract of the brainstem; LL: lateral lemniscus of the brainstem.

**Table 7:** Seed-to-seed functional connectivity (gPPI) for happy > neutral normal > f0 & energy synthesized voices contrast, *p*<.05 seed-level FDR correction, two-tailed.

| 1029 | Seed region label | Target region label | T value | pFDR value |
| --- | --- | --- | --- | --- |
| 1030 | PaCC r | SubCC | -4.73 | 0.0442 |
| 1031 | pMTG l | COC r | 4.70 | 0.0374 |
| 1032 | pMTG l | pSTG r | -4.45 | 0.0374 |
| 1033 | aSTG r | GPI r | 4.70 | 0.0466 |
| 1034 | LL r | CST r | 8.22 | 0.0001 |
| 1035 | PaCC: paracingulate cortex; SubCC: subcalcarine cortex; GPI: internal globus pallidus; COC: central operculum |  |  |  |
| 1036 | cortex; aSTG: anterior superior temporal gyrus; pSTG: posterior superior temporal gyrus; pMTG: posterior |  |  |  |
| 1037 | middle temporal gyrus; CST: corticospinal tract of the brainstem; LL: lateral lemniscus of the brainstem.; l: left; |  |  |  |
| 1038 | r: right. |  |  |  |

### Seed-to-voxel effective connectivity with the basal ganglia as seeds

In order to determine the direct relations between BG regions and the rest of the brain, namely each voxel, we computed seed-to-voxel analyses using multivariate regressions and took as seeds only the BG (N=22 ROI; Fig.6). We only observed significant effective connectivity specific to angry ̶ but not happy ̶ voices through the interaction with the Acoustic Parameters factor [angry > neutral voices * original > *f*0 & energy synthesized voices]. This multivariate analysis revealed a direct coupling between the left STN (seed) and the ipsilateral cerebellum crus II of ansiform lobule (MNI xyz −4 −86 −42; t14=4.14, k=26 voxels; *p*=0.031 FDR corrected, two-tailed; Fig.6A). We also observed an anti-coupling between the left GPe (seed) and left temporo-occipital MTG (MNI xyz −60 −50 −2) and MFG (MNI xyz −44 34 20; for both contrasts, t14=4.14, k= 29 and 20 voxels, respectively; *p*=0.018 and 0.048 FDR corrected, two-tailed, respectively; Fig.6B). Finally, direct coupling was observed between the left caudate nucleus (seed) and voxels covering part of the right primary auditory cortex and planum temporale (MNI xyz 54 −12 0; t14=4.14, k= 64 voxels, *p*=0.00009 FDR corrected, two-tailed; Fig.6C).

**Fig. 6:**
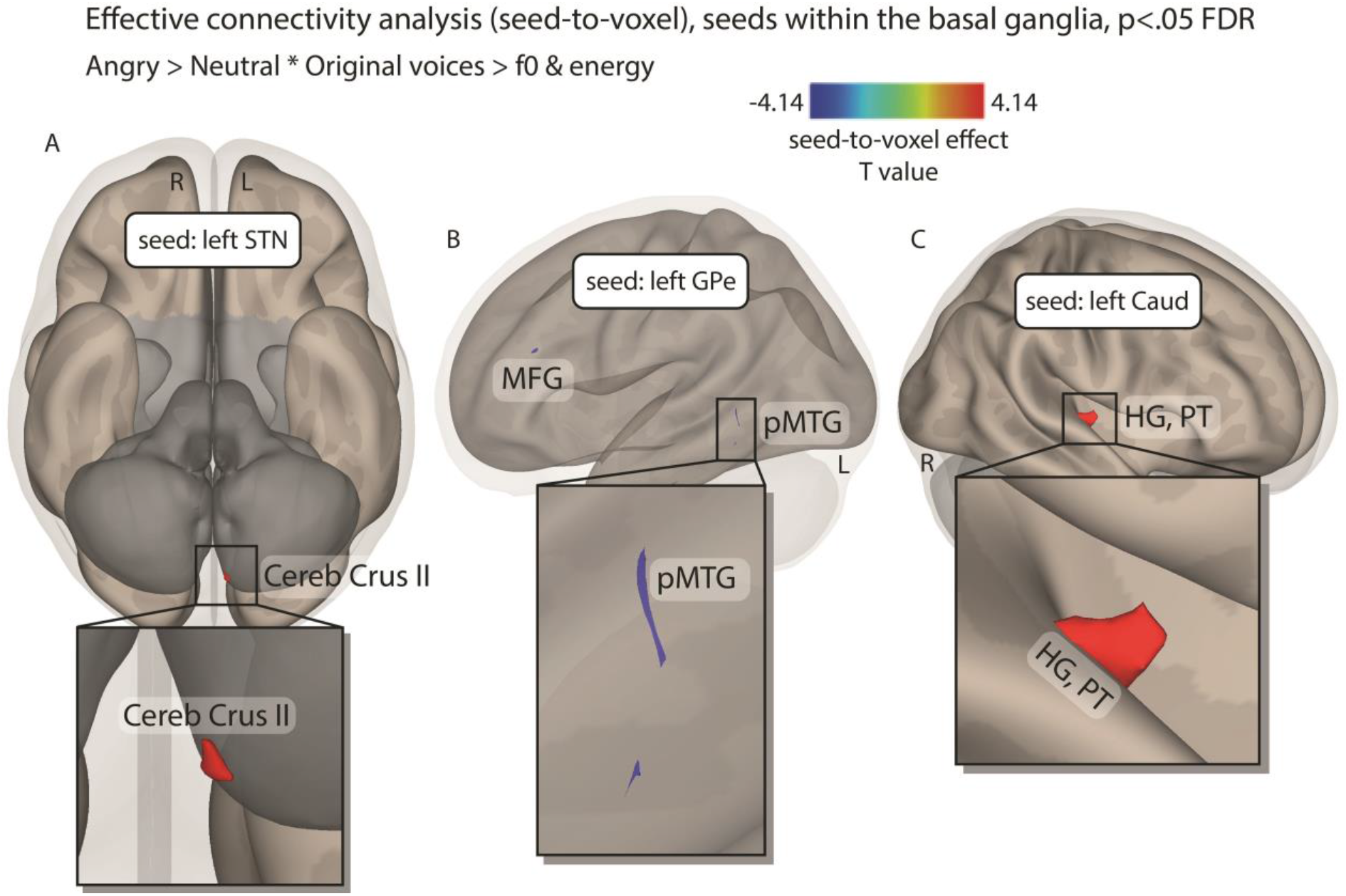
Coupled and anti-coupled seed-to-voxel, gPPI effective connectivity for the interaction between the Emotion and the Acoustic parameter factors contrasting angry > neutral voices * original > *f*0 & energy synthesized voices, corrected for multiple comparisons (*p*<.05 FDR). A, Inferior view showing direct coupling between the left STN (seed) and the ipsilateral Cerebellum Crus II. B, Sagittal view showing direct anti-coupling between the left GPe (seed) and the left MFG and pMTG. C, Sagittal view showing direct coupling between the left caudate nucleus (seed) and the right primary auditory cortex (Heschl’s gyrus) and planum temporale. L: left; R: right; STN: subthalamic nucleus; GPe: external globus pallidus; Caud: caudate nucleus; Cereb Crus II: cerebellum crus II of ansiform lobule; pMTG: posterior middle temporal gyrus; MFG: middle frontal gyrus; HG: Heschl’s gyrus; PT: planum temporale.

### Seed-to-seed effective connectivity within the basal ganglia

We were ultimately interested in the effective connectivity within the basal ganglia when processing emotional (angry, happy) voices and independently of low-level acoustic parameters (synthesized *f*0, energy). We therefore used multiple regression analyses within the BG for our interaction contrasts to highlight direct relations between BG regions. The anger specific contrast [angry > neutral voices * original > *f*0 & energy synthesized voices] did not reveal any effective connectivity in BG regions whereas the happiness specific contrast [happy > neutral voices * original > *f*0 & energy synthesized voices] revealed coupling between the left putamen and GPi (t14=3.78, *p*=0.030 FDR corrected, two-tailed) as well as anti-coupling between the left GPi and the ipsilateral nucleus accumbens (t14=-3.65, *p*=0.039 FDR corrected, two-tailed).

## Discussion

The present study aimed at determining the functional role of both the basal ganglia and cerebellum according to an integrative neural model of vocal emotion perception, decoding and integration using focal, high-resolution fMRI. It was assumed that connectivity – functional and/or effective– between the BG and the cerebellum would underlie the differential processing of emotion, namely angry and/or happy compared to neutral voices, especially when constraining our data by the use of low-level acoustic parameters of no-interest (synthesized *f*0 and synthesized energy voices). Our results confirmed the hypothesized involvement of subparts of the BG and cerebellum in processing vocal emotions. The interaction between emotion and acoustical parameters yielded significant results only for connectivity analyses. Functional connectivity data revealed coupled and anti-coupled networks involving the BG and cerebellum, while effective connectivity within the BG and with the BG as seeds, shed new light on the involvement of the internal and external globus pallidus, putamen, left STN and caudate nucleus in vocal emotion processing.

The implication of subcortical structures other than the amygdala involved in emotion processing was only recently emphasized (Wager, Barrett et al. 2008, Tamietto and De Gelder 2010) and through deep brain stimulation in the STN as a neurosurgical treatment for Parkinson’s disease and obsessive-compulsive disorder, a new research window opened (for a review, see Péron, Frühholz et al. 2013). According to Péron and colleagues’ model (2013) and in line with existing literature and our results, the processing of emotion would rely on both the direct (‘hyperdirect pathway’) and indirect coupling between STN subterritories (motor, associative and limbic) and the neocortex, especially the orbitofrontal cortex (OFC) and modality-specific primary and secondary cortices. Indirect coupling would transit from the STN to the OFC through the BG, especially the GPi and GPe, thalamus, substantia nigra and ventral tegmental area, and/or through the amygdala that exhibits some direct connections with the BG as well (Péron, Frühholz et al. 2013). The STN could synchronize oscillations in relevant areas across the brain including the cerebellum to shape cortical learning and facilitate habitual, overlearned processing of familiar stimuli types (Pierce and Péron 2020). Our results fit well with such model and constrain it by adding some nuance to the expected synchronized regions across the brain. In fact, we observed enhanced activity in several subparts of the BG and in different territories of the cerebellum. More specifically, we observed for angry ̶ similarly for happy ̶ voice processing the involvement of the GPe and thalamus as well as of several lobules (IV,V, VI), nuclei (fastigial, dentate) and areas (Vermis area VI) of the cerebellum and posterior, mid and anterior temporal regions within the voice-sensitive areas. GP activity fits with a more accurate recognition of vocal emotion in healthy compared to BG-lesioned patients (Paulmann, Pell et al. 2008), and with a general role of the more dorsal BG for the sequencing and anticipation of acoustic temporal variations (Kotz, Schwartze et al. 2009). The BG would therefore be crucial to detect and classify auditory patterns, subsequently synchronizing activity in other regions for selecting the appropriate response.

The ‘limbic’ cerebellum −predominately the vermis and posterior lobules, present in our wholebrain and connectivity results– then could modulate these cortical oscillations based on prediction error feedback relative to the given context (Booth, Wood et al. 2007, Schmahmann 2019). By continuously monitoring incoming stimuli for deviations from expected emotional structure (e.g., an angry voice), the limbic cerebellum and especially its subparts -in our results, lobules IV-VI, VIII and the Vermis IV and VI– could signal the need for greater attentional control of sensory cortical responses. Cerebellum activity in our results would also fit well with response adaptation and motor control, preparing a response following vocal emotion decoding and processing (for a review see Frühholz, Trost et al. 2016), especially when the voice or sound is perceived as aversive (Zald and Pardo 2002). Input to the limbic cerebellum (Vermis, cerebellar lobules IV-IX, dentate and fastigial nucleus) from OFC or the BG regarding the salience of emotional stimuli would shape internal models about how an emotional response would affect the individual in its current state, and, thus, how the cerebellum modifies limbic responses, especially in the temporal domain (Breska and Ivry 2018). The idea of temporal pattern analysis in the cerebellum has been proposed, especially when patterns are irregular and not rhythmic (Breska and Ivry 2016), which includes vocal emotion and emotional prosody. Specifically, a double dissociation between patients with a BG or cerebellum lesion confirmed that cerebellar lesions alter non-rhythmic – but not rhythmic – temporal prediction while BG lesions showed the opposite pattern (Breska and Ivry 2018). Additionally, misattributions in emotion recognition between surprise and fear correlated with lesions in lobules VIIb, VIII and X of the cerebellum (Thomasson, Saj et al. 2019), regions that overlap with our results for angry and happy voices in both the wholebrain activation and connectivity analyses and are in line with previous evidence of emotional processing within these specific regions (Stoodley and Schmahmann 2009, Stoodley and Schmahmann 2009, Leggio and Olivito 2018). Therefore, these cerebellar lobules may play a crucial function in emotion recognition in voices, notably in temporal pattern analysis and critical low-level acoustics integration such as *f*0 or pitch.

The importance of BG-cerebellum connections in vocal emotion processing, especially for anger, was further emphasized by our functional connectivity data for angry, but not happy, original voice processing (removing the variance explained by synthesized *f*0 and energy), which revealed coupling between the GPi and putamen with lobule X of the cerebellum. These results are consistent with a coupling of BG and cerebellum activity in time for autonomic emotional reaction and prediction generation (Annoni, Ptak et al. 2003) but cerebellar lobule X is more rarely observed in emotion-related tasks. This cerebellar lobule was however recently integrated in the ‘triple nonmotor representation’ and evidence shows its limbic ties with the neocortex (Guell, Schmahmann et al. 2018). It is also important to note here that many cerebellar sub-regions often labelled as ‘motor’ (for example, linked to hand or eye movements) are also significantly involved in cognitive or emotional tasks (Stoodley and Schmahmann 2010, Stoodley, Valera et al. 2012), a good example concerns lobules V, VI, VIII (King, Hernandez-Castillo et al. 2019). Our results therefore converge toward a critical role of the cerebellum in coordination with the BG for both the decoding of vocal emotion ̶ in the temporal, voice-sensitive areas ̶ and the conversion to a motor response as an output behaviour following a subjective feeling of emotion (Frühholz, Trost et al. 2016, Pierce and Péron 2020).

Furthermore, our effective connectivity results strongly emphasized within-BG direct relations between the putamen and GPi (coupling) and between the GPi and nucleus accumbens (anti-coupling) as well as between BG seeds and frontal and superior temporal regions. Additionally, effective seed-to-voxel connectivity revealed direct coupling between the left STN and ipsilateral cerebellum crus II of the ansiform lobule. While the role of the STN in emotion processing (Schneider, Habel et al. 2003, Kühn, Hariz et al. 2005, Mallet, Schüpbach et al. 2007, Sieger, Serranová et al. 2015, Péron 2016) and vocal emotion recognition (Péron, Grandjean et al. 2010, Péron, Frühholz et al. 2013, Péron, Frühholz et al. 2015, Frühholz, Trost et al. 2016, Péron, Renaud et al. 2017) has gathered strong interest in the recent years, the crus II area of the cerebellum also subserves cognition and emotion processes (Schmahmann 2001, Baumann and Mattingley 2012, Adamaszek, D’Agata et al. 2017). Direct coupling was also observed between the left caudate nucleus and the primary auditory cortex and planum temporale, fitting well again with the direct coupling between the BG and modality-specific sensory cortex (Péron, Frühholz et al. 2013) with the caudate playing a critical role in voice arousal (Bestelmeyer, Kotz et al. 2017) and emotion processing (Grandjean 2017).

We interestingly also observed direct anti-coupling between the left GPe, involved in the explicit recognition of emotional prosody (Paulmann, Pell et al. 2008), and ipsilateral posterior MTG and MFG, superior to and slightly overlapping with the triangularis part of the IFG. Activity modulations in these latter lateral brain areas were repeatedly observed in voice processing in general (Aglieri, Chaminade et al. 2018) and vocal emotion (Leitman, Wolf et al. 2010, Witteman, Van Heuven et al. 2012), especially when contrasting happy to angry voices (Johnstone, Van Reekum et al. 2006). The fact that posterior MTG activity was previously linked to happy vs. angry voice processing therefore could explain the coupling we observed that is specific to happy voices, especially since GP functioning relates to explicit vs. implicit emotion recognition (Paulmann, Pell et al. 2008).

While our data depict a relatively clear image of the importance of the BG and cerebellum for vocal emotion processing and further output response, some limitations should be mentioned. First, sample size was limited and even though we were strict with the correction of *p* values in our statistical analyses, a sample size closer to 25 participants would have been better for reliable data generalization. Second, and as often observed in the literature, we included happy, angry and neutral emotions as vocal stimuli but other critical emotions such as fear, surprise, sadness or several others were not included, therefore restricting our conclusions. Third, although we did include low-level acoustic parameters to control for emotion-specific activity, other meaningful ones should be used in the future, for instance the spectral domain related to voice quality perception, which is thought also important for emotional voice recognition. Fourth, we used high-resolution fMRI, greatly improving spatial resolution with, however, the added cost of a truncated field of view. We cannot therefore exclude the fact that frontal and parietal regions, excluded at data acquisition, would play a role in vocal emotion processing, in terms of both activation and connectivity. It is, however, worth mentioning that the focus of the present study was on cerebellar and basal ganglia contributions to vocal emotion processing. Fifth, we did not divide the STN and other BG or cerebellar regions into their known associative, motor and limbic subparts. A more precise understanding of the specific role of each subpart of the BG nuclei is therefore unfortunately not possible at this stage. Such concern should be addressed in the future by the use of subject-level delineation of BG sub-territories and/or by using even higher fMRI resolution, such as with a 7-tesla scanner. Finally, while our functional connectivity results were consistent with existing literature, we cannot rule out that other regions may mediate the correlations between ROI, so these should be taken with more caution than the effective connectivity results that used more direct mathematical association calculations (multiple regressions). In addition to these limitations, future studies should try to highlight emotional substrates within the BG and cerebellum pertaining to sub-components of emotion, such as for example perception and/or decoding, subjective feeling, response output, behavioural response to emotion, as well as giving more importance to task designs allowing for a clearer topography and parcellation of the affective BG and cerebellum.

In conclusion, the present study aimed at a better understanding of the implications of basal ganglia and cerebellum involvement in vocal emotion processing. Through the combination of wholebrain analysis, functional and effective connectivity analyses and with the partial exclusion of low-level acoustics of interest (voice *f*0, energy) our data depict a clearer role of the STN, GP and putamen in vocal emotion processing, especially for auditory pattern detection and synchronization across cortical and subcortical limbic networks. The current results add weight to the assertion that both direct and indirect coupling between these BG regions and the cortex is modulated by BG and cerebellum connections. Our results also favour a framework in which the brain could use temporal regularities (‘patterns’) to analyse and anticipate the timing of future events, and constrain attention and action accordingly. Further work use a dedicated task and focus on BG and cerebellum subterritories since their specific role(s) is of the highest interest for affective and social neuroscience research.

## Acknowledgements

The present study was performed at the Brain and Behaviour Laboratory and at the Swiss Center for Affective Sciences of the University of Geneva and was funded by Swiss National Foundation grant no. 105314_ 1406 22 (DG-JP) and 105314_182221 (JP). The funders had no role in data collection, discussion of content, preparation of the manuscript, or decision to publish. We would like to thank the healthy controls for contributing their time to this study.

## References

Adamaszek, M., F. D’Agata, R. Ferrucci, C. Habas, S. Keulen, K. Kirkby, M. Leggio, P. Mariën, M. Molinari and E. Moulton (2017). “Consensus paper: cerebellum and emotion.” The Cerebellum 16(2): 552–576.

Aglieri, V., T. Chaminade, S. Takerkart and P. Belin (2018). “Functional connectivity within the voice perception network and its behavioural relevance.” NeuroImage 183: 356–365.

Alexander, G. E. and M. D. Crutcher (1990). “Functional architecture of basal ganglia circuits: neural substrates of parallel processing.” Trends in neurosciences 13(7): 266–271.

Amunts, K., C. Lepage, L. Borgeat, H. Mohlberg, T. Dickscheid, M.-É. Rousseau, S. Bludau, P.-L. Bazin, L. B. Lewis and A.-M. Oros-Peusquens (2013). “BigBrain: an ultrahigh-resolution 3D human brain model.” Science 340(6139): 1472–1475.

Annoni, J. M., R. Ptak, A. S. Caldara-Schnetzer, A. Khateb and B. Z. Pollermann (2003). “Decoupling of autonomic and cognitive emotional reactions after cerebellar stroke.” Annals of Neurology: Official Journal of the American Neurological Association and the Child Neurology Society 53(5): 654–658.

Ashburner, J. (2007). “A fast diffeomorphic image registration algorithm.” Neuroimage 38(1): 95–113.

Banse, R. and K. R. Scherer (1996). “Acoustic profiles in vocal emotion expression.” J Pers Soc Psychol 70(3): 614–636.

Banziger, T. and K. R. Scherer (2010). Introducing the Geneva Multimodal Emotion Portrayal (GEMEP) Corpus A blueprint for an affectively competent agent: Cross-fertilization between Emotion Psychology, Affective Neuroscience, and Affective Computing. T. Banziger, K. Scherer and E. Roesch. Oxford, Oxford University Press.

Baumann, O. and J. B. Mattingley (2012). “Functional topography of primary emotion processing in the human cerebellum.” NeuroImage 61(4): 805–811.

Bestelmeyer, P. E., S. A. Kotz and P. Belin (2017). “Effects of emotional valence and arousal on the voice perception network.” Social cognitive and affective neuroscience 12(8): 1351–1358.

Booth, J. R., L. Wood, D. Lu, J. C. Houk and T. Bitan (2007). “The role of the basal ganglia and cerebellum in language processing.” Brain Research 1133: 136–144.

Bostan, A. C. and P. L. Strick (2018). “The basal ganglia and the cerebellum: nodes in an integrated network.” Nat Rev Neurosci 19(6): 338–350.

Breska, A. and R. B. Ivry (2016). “Taxonomies of timing: where does the cerebellum fit in?” Current opinion in behavioral sciences 8: 282–288.

Breska, A. and R. B. Ivry (2018). “Double dissociation of single-interval and rhythmic temporal prediction in cerebellar degeneration and Parkinson’s disease.” Proceedings of the National Academy of Sciences 115(48): 12283–12288.

Buckner, R. L., F. M. Krienen, A. Castellanos, J. C. Diaz and B. T. Yeo (2011). “The organization of the human cerebellum estimated by intrinsic functional connectivity.” Journal of neurophysiology.

Cohen, M. J., C. A. Riccio and A. M. Flannery (1994). “Expressive aprosodia following stroke to the right basal ganglia: A case report.” Neuropsychology 8(2): 242.

Collins, D. L., P. Neelin, T. M. Peters and A. C. Evans (1994). “Automatic 3D intersubject registration of MR volumetric data in standardized Talairach space.” Journal of computer assisted tomography 18(2): 192–205.

Diedrichsen, J., J. H. Balsters, J. Flavell, E. Cussans and N. Ramnani (2009). “A probabilistic MR atlas of the human cerebellum.” Neuroimage 46(1): 39–46.

Diedrichsen, J., S. Maderwald, M. Küper, M. Thürling, K. Rabe, E. Gizewski, M. E. Ladd and D. Timmann (2011). “Imaging the deep cerebellar nuclei: a probabilistic atlas and normalization procedure.” Neuroimage 54(3): 1786–1794.

Ethofer, T., S. Anders, M. Erb, C. Herbert, S. Wiethoff, J. Kissler, W. Grodd and D. Wildgruber (2006). “Cerebral pathways in processing of affective prosody: a dynamic causal modeling study.” Neuroimage 30(2): 580–587.

Fonov, V., A. C. Evans, K. Botteron, C. R. Almli, R. C. McKinstry, D. L. Collins and B. D. C. Group (2011). “Unbiased average age-appropriate atlases for pediatric studies.” Neuroimage 54(1): 313–327.

Frühholz, S., L. Ceravolo and D. Grandjean (2011). “Specific brain networks during explicit and implicit decoding of emotional prosody.” Cerebral cortex 22(5): 1107–1117.

Frühholz, S., L. Ceravolo and D. Grandjean (2012). “Specific brain networks during explicit and implicit decoding of emotional prosody.” Cerebral cortex 22(5): 1107–1117.

Frühholz, S. and D. Grandjean (2013). “Multiple subregions in superior temporal cortex are differentially sensitive to vocal expressions: a quantitative meta-analysis.” Neuroscience & Biobehavioral Reviews 37(1): 24–35.

Frühholz, S. and D. Grandjean (2013). “Processing of emotional vocalizations in bilateral inferior frontal cortex.” Neuroscience & Biobehavioral Reviews 37(10): 2847–2855.

Frühholz, S., W. Trost and S. A. Kotz (2016). “The sound of emotions—Towards a unifying neural network perspective of affective sound processing.” Neuroscience & Biobehavioral Reviews 68: 96–110.

Grandjean, D. (2017). Brain Mechanisms in Emotional Voice Production and Perception and Early Life Interactions. Early Vocal Contact and Preterm Infant Brain Development, Springer :71–87.

Grandjean, D. (in press). “Brain networks of emotional prosody processing.” Emotion Review.

Grandjean, D., T. Banziger and K. R. Scherer (2006). “Intonation as an interface between language and affect.” Prog Brain Res 156: 235–247.

Graybiel, A. M. (2008). “Habits, rituals, and the evaluative brain.” Annu Rev Neurosci 31: 359–387.

Grube, M., F. E. Cooper, P. F. Chinnery and T. D. Griffiths (2010). “Dissociation of duration-based and beat-based auditory timing in cerebellar degeneration.” Proceedings of the National Academy of Sciences 107(25): 11597–11601.

Guell, X., J. D. Schmahmann, J. D. Gabrieli and S. S. Ghosh (2018). “Functional gradients of the cerebellum.” Elife 7: e36652.

Habas, C., N. Kamdar, D. Nguyen, K. Prater, C. F. Beckmann, V. Menon and M. D. Greicius (2009). “Distinct cerebellar contributions to intrinsic connectivity networks.” Journal of neuroscience 29(26): 8586–8594.

Johnstone, T., C. M. Van Reekum, T. R. Oakes and R. J. Davidson (2006). “The voice of emotion: an FMRI study of neural responses to angry and happy vocal expressions.” Social Cognitive and Affective Neuroscience 1(3): 242–249.

King, M., C. R. Hernandez-Castillo, R. A. Poldrack, R. B. Ivry and J. Diedrichsen (2019). “Functional boundaries in the human cerebellum revealed by a multi-domain task battery.” Nature neuroscience 22(8): 1371–1378.

Kotz, S. A. and M. Schwartze (2010). “Cortical speech processing unplugged: a timely subcortico-cortical framework.” Trends in cognitive sciences 14(9): 392–399.

Kotz, S. A., M. Schwartze and M. Schmidt-Kassow (2009). “Non-motor basal ganglia functions: A review and proposal for a model of sensory predictability in auditory language perception.” Cortex 45(8): 982–990.

Kühn, A., M. Hariz, P. Silberstein, S. Tisch, A. Kupsch, G.-H. Schneider, P. Limousin-Dowsey, K. Yarrow and P. Brown (2005). “Activation of the subthalamic region during emotional processing in Parkinson disease.” Neurology 65(5): 707–713.

Lambert, C., L. Zrinzo, Z. Nagy, A. Lutti, M. Hariz, T. Foltynie, B. Draganski, J. Ashburner and R. Frackowiak (2012). “Confirmation of functional zones within the human subthalamic nucleus: patterns of connectivity and sub-parcellation using diffusion weighted imaging.” Neuroimage 60(1): 83–94.

Larry, N., M. Yarkoni, A. Lixenberg and M. Joshua (2019). “Cerebellar climbing fibers encode expected reward size.” bioRxiv: 533653.

Leggio, M. and G. Olivito (2018). Topography of the cerebellum in relation to social brain regions and emotions. Handbook of clinical neurology, Elsevier. 154:71–84.

Leitman, D. I., D. H. Wolf, J. D. Ragland, P. Laukka, J. Loughead, J. N. Valdez, D. C. Javitt, B. Turetsky and R. Gur (2010). ““ It’s not what you say, but how you say it”: a reciprocal temporo-frontal network for affective prosody.” Frontiers in human neuroscience 4: 19.

Mallet, L., M. Schüpbach, K. N’Diaye, P. Remy, E. Bardinet, V. Czernecki, M.-L. Welter, A. Pelissolo, M. Ruberg and Y. Agid (2007). “Stimulation of subterritories of the subthalamic nucleus reveals its role in the integration of the emotional and motor aspects of behavior.” Proceedings of the National Academy of Sciences 104(25): 10661–10666.

Paulmann, S., M. D. Pell and S. A. Kotz (2008). “Functional contributions of the basal ganglia to emotional prosody: Evidence from ERPs.” Brain Research 1217: 171–178.

Pell, M. D. and C. L. Leonard (2003). “Processing emotional tone from speech in Parkinson’s disease: a role for the basal ganglia.” Cognitive, Affective, & Behavioral Neuroscience 3(4): 275–288.

Péron, J. (2016). “The role of the subthalamic nucleus in emotional processing.” Clinical Neurophysiology 127(3): e39.

Péron, J., S. Frühholz, L. Ceravolo and D. Grandjean (2015). “Structural and functional connectivity of the subthalamic nucleus during vocal emotion decoding.” Social cognitive and affective neuroscience 11(2): 349–356.

Péron, J., S. Frühholz, L. Ceravolo and D. Grandjean (2016). “Structural and functional connectivity of the subthalamic nucleus during vocal emotion decoding.” Soc Cogn Affect Neurosci 11(2): 349–356.

Péron, J., S. Frühholz, M. Vérin and D. Grandjean (2013). “Subthalamic nucleus: A key structure for emotional component synchronization in humans.” Neurosci Biobehav Rev 37(3): 358–373.

Péron, J., S. Frühholz, M. Vérin and D. Grandjean (2013). “Subthalamic nucleus: A key structure for emotional component synchronization in humans.” Neuroscience & Biobehavioral Reviews 37(3): 358–373.

Péron, J., D. Grandjean, F. Le Jeune, P. Sauleau, C. Haegelen, D. Drapier, T. Rouaud, S. Drapier and M. Vérin (2010). “Recognition of emotional prosody is altered after subthalamic nucleus deep brain stimulation in Parkinson’s disease.” Neuropsychologia 48(4): 1053–1062.

Péron, J., C. Haegelen, P. Sauleau, L. Tamarit, V. Milesi, J. F. Houvenaghel, T. Dondaine, M. Vérin and D. Grandjean (2014). “Electrophysiological activity of the subthalamic nucleus in response to emotional prosody: An intracranial ERP study in Parkinson’s disease.” Movement Disorders 29(S284-S285).

Péron, J., O. Renaud, C. Haegelen, L. Tamarit, V. Milesi, J.-F. Houvenaghel, T. Dondaine, M. Vérin, P. Sauleau and D. Grandjean (2017). “Vocal emotion decoding in the subthalamic nucleus: An intracranial ERP study in Parkinson’s disease.” Brain and language 168: 1–11.

Pierce, J. E. and J. Péron (2020). “The basal ganglia and the cerebellum in human emotion.” Social Cognitive and Affective Neuroscience.

Reid, A. T., D. B. Headley, R. D. Mill, R. Sanchez-Romero, L. Q. Uddin, D. Marinazzo, D. J. Lurie, P. A. Valdés-Sosa, S. J. Hanson and B. B. Biswal (2019). “Advancing functional connectivity research from association to causation.” Nature neuroscience 1(10).

Schirmer, A. and S. A. Kotz (2006). “Beyond the right hemisphere: brain mechanisms mediating vocal emotional processing.” Trends in cognitive sciences 10(1): 24–30.

Schmahmann, J. D. (2001). “The cerebrocerebellar system: Anatomic substrates of the cerebellar contribution to cognition and emotion.” International Review of Psychiatry 13(4): 247–260.

Schmahmann, J. D. (2019). “The cerebellum and cognition.” Neurosci Lett 688: 62–75.

Schneider, F., U. Habel, J. Volkmann, S. Regel, J. Kornischka, V. Sturm and H.-J. Freund (2003). “Deep brain stimulation of the subthalamic nucleus enhances emotional processing in Parkinson disease.” Archives of general psychiatry 60(3): 296–302.

Sieger, T., T. Serranová, F. Růžička, P. Vostatek, J. Wild, D. Šťastná, C. Bonnet, D. Novák, E. Růžička and D. Urgošík (2015). “Distinct populations of neurons respond to emotional valence and arousal in the human subthalamic nucleus.” Proceedings of the National Academy of Sciences 112(10): 3116–3121.

Smith, S. M., M. Jenkinson, M. W. Woolrich, C. F. Beckmann, T. E. Behrens, H. Johansen-Berg, P. R. Bannister, M. De Luca, I. Drobnjak and D. E. Flitney (2004). “Advances in functional and structural MR image analysis and implementation as FSL.” Neuroimage 23: S208–S219.

Sokolov, A. A., R. C. Miall and R. B. Ivry (2017). “The Cerebellum: Adaptive Prediction for Movement and Cognition.” Trends Cogn Sci 21(5): 313–332.

Stoodley, C. J. and J. D. Schmahmann (2009). “The cerebellum and language: evidence from patients with cerebellar degeneration.” Brain and language 110(3): 149–153.

Stoodley, C. J. and J. D. Schmahmann (2009). “Functional topography in the human cerebellum: a meta-analysis of neuroimaging studies.” Neuroimage 44(2): 489–501.

Stoodley, C. J. and J. D. Schmahmann (2010). “Evidence for topographic organization in the cerebellum of motor control versus cognitive and affective processing.” Cortex 46(7): 831–844.

Stoodley, C. J., E. M. Valera and J. D. Schmahmann (2012). “Functional topography of the cerebellum for motor and cognitive tasks: an fMRI study.” Neuroimage 59(2): 1560–1570.

Tamietto, M. and B. De Gelder (2010). “Neural bases of the non-conscious perception of emotional signals.” Nature Reviews Neuroscience 11(10): 697.

Thomasson, M., A. Saj, D. Benis, D. Grandjean, F. Assal and J. Péron (2019). “Cerebellar contribution to vocal emotion decoding: Insights from stroke and neuroimaging.” Neuropsychologia 132: 107141.

Tzourio-Mazoyer, N., B. Landeau, D. Papathanassiou, F. Crivello, O. Etard, N. Delcroix, B. Mazoyer and M. Joliot (2002). “Automated anatomical labeling of activations in SPM using a macroscopic anatomical parcellation of the MNI MRI single-subject brain.” Neuroimage 15(1): 273–289.

Wager, T. D., L. F. Barrett, E. Bliss-Moreau, K. Lindquist, S. Duncan, H. Kober, J. Joseph, M. Davidson and J. Mize (2008). “The neuroimaging of emotion.” Handbook of emotions 3: 249–271.

Wang, J., W. W. Dong, W. H. Zhang, J. Zheng and X. Wang (2014). “Electrical stimulation of cerebellar fastigial nucleus: mechanism of neuroprotection and prospects for clinical application against cerebral ischemia.” CNS neuroscience & therapeutics 20(8): 710–716.

Whitfield-Gabrieli, S. and A. Nieto-Castanon (2012). “Conn: a functional connectivity toolbox for correlated and anticorrelated brain networks.” Brain connectivity 2(3): 125–141.

Witteman, J., V. J. Van Heuven and N. O. Schiller (2012). “Hearing feelings: a quantitative meta-analysis on the neuroimaging literature of emotional prosody perception.” Neuropsychologia 50(12): 2752–2763.

Zald, D. H. and J. V. Pardo (2002). “The neural correlates of aversive auditory stimulation.” Neuroimage 16(3): 746–753.

Zhang, X.-Y., J.-J. Wang and J.-N. Zhu (2016). “Cerebellar fastigial nucleus: from anatomic construction to physiological functions.” Cerebellum & Ataxias 3(1): 9.

