## Supplementary figures and tables for "Basal ganglia and cerebellar contributions to vocal emotion processing: a high resolution fMRI study"

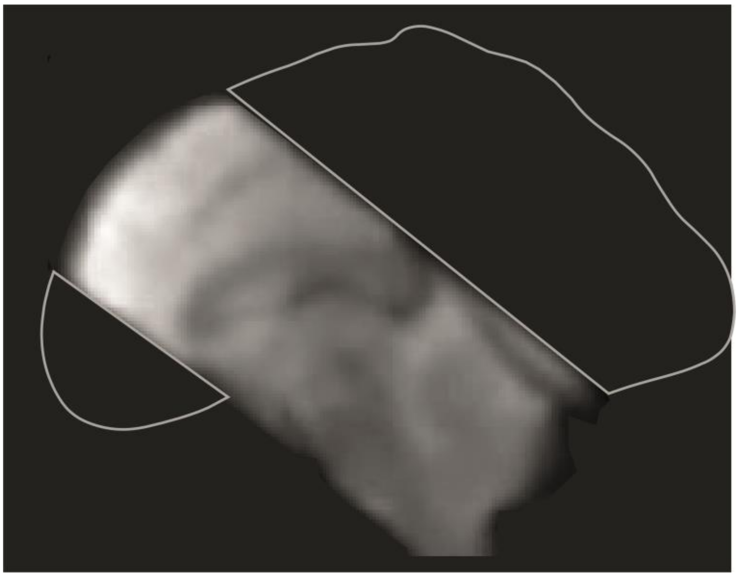

Fig.S1: Illustration of the truncated field of view of the high-resolution functional MRI scans acquired continuously during the one-back task.

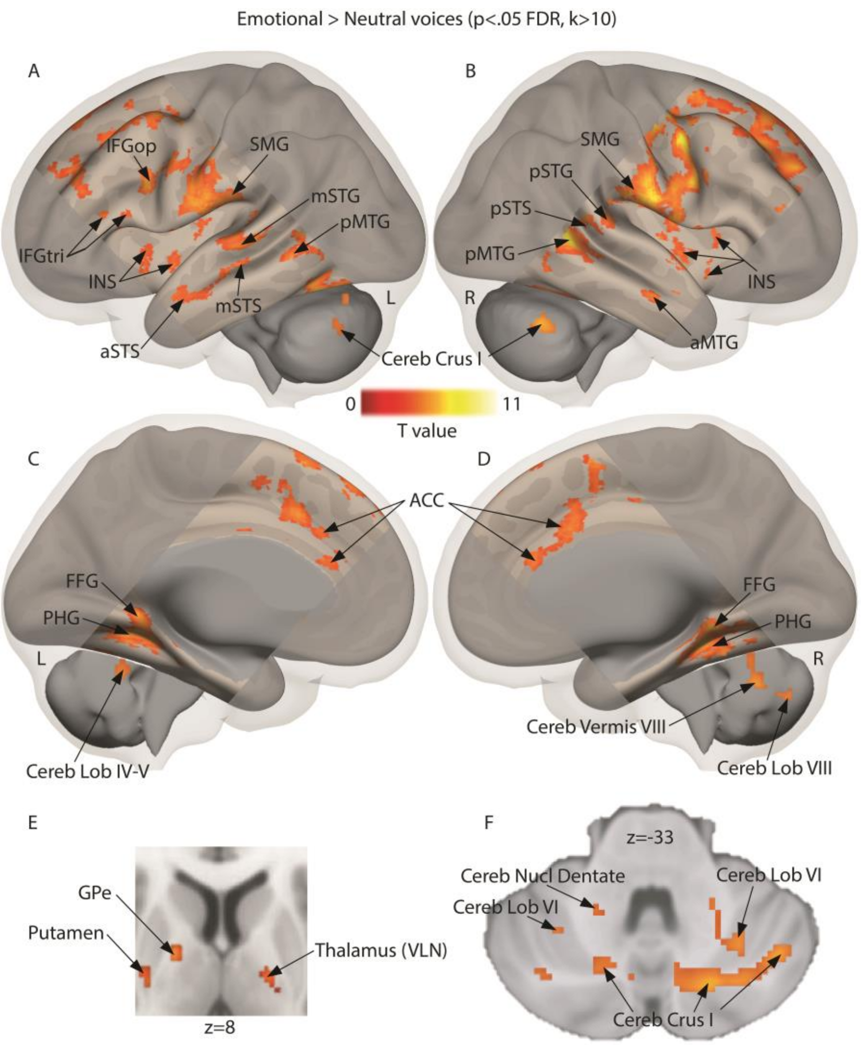

Fig.S2: Brain measures for implicitly processing angry and happy compared to neutral voices, corrected for multiple comparisons (whole brain voxelwise *p*<.05 FDR, k>10 voxels). A-B Lateral activations rendered on a sagittal image highlighting middle and superior temporal regions. C-D Medial activations of the anterior cingulate cortex, parahippocampal cortex and cerebellum. E Subcortical activity in the thalamus and globus pallidus displayed on an axial slice. F Cerebellar activations displayed on an axial slice. White outline: sample-specific temporal voice areas, thresholded at whole brain voxelwise *p*<.05 FDR corrected at the voxel level. L: left; R: right; IFGop: inferior frontal gyrus pars opercularis; IFGtri: inferior frontal gyrus pars triangularis; STG: superior temporal gyrus; STS: superior temporal sulcus; MTG: middle temporal gyrus; STS: superior temporal sulcus; INS: insula; SMG: supramarginal gyrus; FG: frontal gyrus; FFG: fusiform gyrus; PHG: parahippocampal gyrus; ACC: anterior cingulate cortex; Cereb: cerebellum; Cereb Lob: cerebellum lobule; Cereb Nucl Dentate: dentate nucleus of the cerebellum; Thalamus VLN: ventral lateral nucleus of the thalamus;

GPe: external globus pallidus; Cereb Crus: cerebellum crus of ansiform lobule. ‘a’ prefix: anterior part; ‘m’ prefix: mid part; ‘p’ prefix: posterior part.

Supplementary Table 1: Regions of interest included in the functional and/or effective connectivity analyses, all within the field of view of the acquired MRI volumes.

aal.IC r (Insular Cortex Right)

aal.IC l (Insular Cortex Left)

aal.SFG r (Superior Frontal Gyrus Right)

aal.SFG l (Superior Frontal Gyrus Left)

aal.MidFG r (Middle Frontal Gyrus Right)

aal.MidFG l (Middle Frontal Gyrus Left)

aal.PreCG r (Precentral Gyrus Right)

aal.PreCG l (Precentral Gyrus Left)

aal.aSTG r (Superior Temporal Gyrus, anterior division Right)

aal.aSTG l (Superior Temporal Gyrus, anterior division Left)

aal.pSTG r (Superior Temporal Gyrus, posterior division Right)

aal.pSTG l (Superior Temporal Gyrus, posterior division Left)

aal.aMTG r (Middle Temporal Gyrus, anterior division Right)

aal.aMTG l (Middle Temporal Gyrus, anterior division Left)

aal.pMTG r (Middle Temporal Gyrus, posterior division Right)

aal.pMTG l (Middle Temporal Gyrus, posterior division Left)

aal.toMTG r (Middle Temporal Gyrus, temporooccipital part Right)

aal.toMTG l (Middle Temporal Gyrus, temporooccipital part Left)

aal.aITG r (Inferior Temporal Gyrus, anterior division Right)

aal.aITG l (Inferior Temporal Gyrus, anterior division Left)

aal.pITG r (Inferior Temporal Gyrus, posterior division Right)

aal.pITG l (Inferior Temporal Gyrus, posterior division Left)

aal.toITG r (Inferior Temporal Gyrus, temporooccipital part Right)

aal.toITG l (Inferior Temporal Gyrus, temporooccipital part Left)

aal.PostCG r (Postcentral Gyrus Right)

aal.PostCG l (Postcentral Gyrus Left)

aal.aSMG r (Supramarginal Gyrus, anterior division Right)

aal.aSMG l (Supramarginal Gyrus, anterior division Left)

aal.pSMG r (Supramarginal Gyrus, posterior division Right)

aal.pSMG l (Supramarginal Gyrus, posterior division Left)

aal.AG r (Angular Gyrus Right)

aal.AG l (Angular Gyrus Left)

aal.SMA r (Juxtapositional Lobule Cortex - formerly Supplementary Motor Cortex- Right)

aal.SMA L(Juxtapositional Lobule Cortex - formerly Supplementary Motor Cortex- Left)

aal.SubCalC (Subcallosal Cortex)

aal.PaCiG r (Paracingulate Gyrus Right)

aal.PaCiG l (Paracingulate Gyrus Left)

aal.PC (Cingulate Gyrus, posterior division)

aal.aPaHC r (Parahippocampal Gyrus, anterior division Right)

aal.aPaHC l (Parahippocampal Gyrus, anterior division Left)

aal.pPaHC r (Parahippocampal Gyrus, posterior division Right)

aal.pPaHC l (Parahippocampal Gyrus, posterior division Left)

aal.aTFusC r (Temporal Fusiform Cortex, anterior division Right)

aal.aTFusC l (Temporal Fusiform Cortex, anterior division Left)

aal.pTFusC r (Temporal Fusiform Cortex, posterior division Right)

aal.pTFusC l (Temporal Fusiform Cortex, posterior division Left)

aal.FO r (Frontal Operculum Cortex Right)
aal.FO l (Frontal Operculum Cortex Left)
aal.CO r (Central Opercular Cortex Right)
aal.CO l (Central Opercular Cortex Left)
aal.PO r (Parietal Operculum Cortex Right)
aal.PO l (Parietal Operculum Cortex Left)
aal.PP r (Planum Polare Right)
aal.PP l (Planum Polare Left)
aal.HG r (Heschl's Gyrus Right)
aal.HG l (Heschl's Gyrus Left)
aal.PT r (Planum Temporale Right)
aal.PT l (Planum Temporale Left)
BG.1 left red nucleus
BG.2 right red nucleus
BG.3 left substantia nigra
BG.4 right substantia nigra
BG.5 left subthalamic nucleus
BG.6 right subthalamic nucleus
BG.7 left caudate
BG.8 right caudate
BG.9 left putamen
BG.10 right putamen
BG.11 left external globus pallidus
BG.12 right external globus pallidus
BG.13 left internal globus pallidus
BG.14 right internal globus pallidus
BG.15 left thalamus
BG.16 right thalamus
BG.17 left hippocampus
BG.18 right hippocampus
BG.19 left nucleus accumbens
BG.20 right nucleus accumbens
BG.21 left amygdala
BG.22 right amygdala

Cerebellum.1 Left_I_IV
Cerebellum.2 Right_I_IV
Cerebellum.3 Left_V
Cerebellum.4 Right_V
Cerebellum.5 Left_VI
Cerebellum.6 Vermis_VI
Cerebellum.7 Right_VI
Cerebellum.8 Left_CrusI
Cerebellum.9 Vermis_CrusI
Cerebellum.10 Right_CrusI
Cerebellum.11 Left_CrusII
Cerebellum.12 Vermis_CrusII
Cerebellum.13 Right_CrusII
Cerebellum.14 Left_VIIb
Cerebellum.15 Vermis_VIIb
Cerebellum.16 Right_VIIb
Cerebellum.17 Left_VIIIa
Cerebellum.18 Vermis_VIIIa
Cerebellum.19 Right_VIIIa
Cerebellum.20 Left_VIIIb
Cerebellum.21 Vermis_VIIIb
Cerebellum.22 Right_VIIIb
Cerebellum.23 Left_IX
Cerebellum.24 Vermis_IX
Cerebellum.25 Right_IX
Cerebellum.26 Left_X
Cerebellum.27 Vermis_X
Cerebellum.28 Right_X
Cerebellum.29 Left_Dentate nucleus
Cerebellum.30 Right_Dentate nucleus
Cerebellum.31 Left_Interposed nucleus
Cerebellum.32 Right_Interposed nucleus
Cerebellum.33 Left_Fastigial nucleus
Cerebellum.34 Right_Fastigial nucleus

Brainstem.CST Left (Corticospinal; motor)

Brainstem.CST Right (Corticospinal; motor)

Brainstem.FPT Left (Fronto-pontine; motor)

Brainstem.FPT Right (Fronto-pontine; motor)

Brainstem.ICPMC Left (Inferior Cerebellar ; cerebellar peduncles)

Brainstem.ICPMC Right (Inferior Cerebellar ; cerebellar peduncles)

Brainstem.ICPVC Left (Inferior Cerebellar; cerebellar peduncles)

Brainstem.ICPVC Right (Inferior Cerebellar; cerebellar peduncles)

Brainstem.LL Left (Lateral lemniscus; sensory tracts)

Brainstem.LL Right (Lateral lemniscus; sensory tracts)

Brainstem.MCP (Middle Cerebellar; cerebellar peduncles)

Brainstem.ML Left (Medial lemniscus; sensory tracts)

Brainstem.ML Right (Medial lemniscus; sensory tracts)

Brainstem.POTPT Left (Parieto-occipito-temporal; motor tracts)

Brainstem.POTPT Right (Parieto-occipito-temporal; motor tracts)

Brainstem.SCPCR Left (Superior Cerebellar; cerebellar peduncles)

Brainstem.SCPCR Right (Superior Cerebellar; cerebellar peduncles)

Brainstem.SCPCT Left (Superior Cerebellar; cerebellar peduncles)

Brainstem.SCPCT Right (Superior Cerebellar; cerebellar peduncles)

Brainstem.SCPSC Left (Superior Cerebellar; cerebellar peduncles)

Brainstem.SCPSC Right (Superior Cerebellar; cerebellar peduncles)

Brainstem.STT Left (Spinothalamic; sensory tracts)

Brainstem.STT (Spinothalamic; sensory tracts)

‘aal.’: Automated Anatomical Labelling atlas ($N_{ROI}$=58); ‘BG.’: Basal Ganglia atlas ($N_{ROI}$=22); ‘Cerebellum.’: Cerebellum atlas ($N_{ROI}$=34); ‘Brainstem.’: Brainstem atlas ($N_{ROI}$=23). For references, see Methods section.

Supplementary Table 2: Activations, cluster size and coordinates for normal voices>F0 flattened voices & voice mean energy contrast, whole brain voxelwise $p<.05$ FDR correction, k>10.

| | | MNI coordinates | | | | |
|---|---|---|---|---|---|---|
| Region label | Hemisphere | X | Y | Z | T value | Cluster size (voxels) |
| STG, posterior | R | 54 | -34 | 12 | 14.27 | 7688 |
| *Precentral gyrus* | *R* | *60* | *8* | *26* | *13.35* | |
| ITG, posterior | L | -46 | -36 | -16 | 9.43 | 198 |
| *ITG, posterior* | *L* | *-44* | *-54* | *-12* | *4.29* | |
| *ITG, posterior* | *L* | *-52* | *-44* | *-16* | *4.29* | |
| Cereb Lob 6 | R | 20 | -56 | -16 | 7.93 | 356 |
| *Fusiform gyrus* | *R* | *34* | *-62* | *-20* | *6.82* | |
| *Cereb Lob 6* | *R* | *30* | *-60* | *-28* | *6.10* | |
| Mid Frontal gyrus | R | 36 | 26 | 20 | 7.23 | 994 |
| *Mid Frontal gyrus* | *R* | *24* | *28* | *36* | *7.12* | |
| *Mid Frontal gyrus* | *R* | *28* | *38* | *28* | *7.01* | |
| STS, posterior | R | 60 | -46 | 4 | 7.12 | 37 |
| MTG, anterior | R | 66 | -4 | -16 | 6.90 | 60 |
| Mid Frontal gyrus | L | -38 | 34 | 20 | 6.73 | 376 |
| *IFG triangularis* | *L* | *-52* | *30* | *12* | *5.09* | |
| *Mid Frontal gyrus* | *L* | *-32* | *38* | *24* | *5.05* | |
| IFG opercularis | L | -50 | 10 | 8 | 6.45 | 53 |
| *IFG opercularis* | *L* | *-58* | *14* | *8* | *4.69* | |
| Supp Motor tracts Area | R | 10 | 6 | 58 | 5.87 | 34 |
| Hippocampus, anterior | L | -26 | -6 | -24 | 5.37 | 31 |
| Supp Motor tracts Area | L | -2 | 22 | 60 | 5.21 | 169 |
| Cingulate cortex | R | 10 | 14 | 42 | 5.18 | 43 |
| Cereb Lob 8 | R | 20 | -70 | -44 | 5.11 | 64 |
| Cereb Crus 2 | R | 8 | -82 | -36 | 5.05 | 24 |
| Hippocampus, anterior | L | -12 | -10 | -22 | 4.92 | 38 |
| Cereb Lob 8 | L | -18 | -58 | -20 | 4.87 | 135 |
| Cereb Nucl Dentate | R | 14 | -48 | -28 | 4.75 | 14 |
| STG, mid | R | 66 | -16 | 6 | 4.72 | 22 |
| IFG, triangularis | L | -42 | 18 | 6 | 4.68 | 37 |
| Sup Frontal gyrus, medial | R | 12 | 50 | 34 | 4.61 | 42 |
| Cereb Crus 1 | L | -38 | -80 | -30 | 4.52 | 14 |
| Putamen | L | -16 | 8 | -12 | 4.43 | 19 |
| Cereb Lob 8 | L | -28 | -58 | -52 | 4.43 | 20 |
| Cereb Vermis 4-5 | L | -2 | -58 | -26 | 4.35 | 47 |

| | | | | | | |
|---|---|---|---|---|---|---|
| Sup Frontal gyrus | L | -14 | 36 | 48 | 4.25 | 50 |
| ACC | R | 6 | 16 | 28 | 4.23 | 26 |
| Sup Frontal gyrus, medial | L | -8 | 48 | 34 | 4.22 | 38 |
| ACC | R | 2 | 34 | 20 | 4.05 | 39 |
| Thalamus | L | -18 | -18 | 12 | 4.02 | 37 |
| Globus Pallidus | R | 18 | -4 | 10 | 2.96 | 11 |

L: left; R: right; MNI: Montreal neurological institute; STG: superior temporal gyrus; ITG: inferior temporal gyrus; Cereb Lob: cerebellum lobule; Mid: middle; STS: superior temporal sulcus; MTG: middle temporal gyrus; IFG: inferior frontal gyrus; Supp Motor tracts Area: supplementary motor tracts area; Cereb Crus: cerebellum crus of ansiform lobule; Cereb Nucl Dentate: dentate nucleus of the cerebellum; Sup: superior; Cereb: cerebellum; ACC: anterior cingulate cortex.

Supplementary Table 3: Activations, cluster size and coordinates for F0 flattened voices>normal voices & voice mean energy contrast, whole brain voxelwise *p*<.05 FDR correction, k>10.

| | | MNI coordinates | | | | |
|---|---|---|---|---|---|---|
| Region label | Hemisphere | X | Y | Z | T value | Cluster size (voxels) |
| STG, anterior | L | -62 | -6 | 4 | 19.03 | 12010 |
| *IFG triangularis* | *R* | *58* | *22* | *6* | *16.42* | |
| *STG, posterior* | *L* | *-40* | *-24* | *2* | *15.51* | |
| *IFG opercularis* | *L* | *-60* | *6* | *14* | *13.99* | |
| *IFG opercularis* | *R* | *64* | *6* | *14* | *11.92* | |
| *STG, mid* | *L* | *-48* | *-18* | *2* | *11.91* | |
| *MTG, anterior* | *R* | *60* | *-10* | *-12* | *11.88* | |
| *Precentral gyrus* | *L* | *-62* | *4* | *20* | *11.38* | |
| *STS, mid* | *R* | *60* | *-20* | *-6* | *11.09* | |
| *MTG, anterior* | *R* | *58* | *-6* | *-14* | *10.55* | |
| Supp Motor tracts Area | R | 8 | 14 | 52 | 14.98 | 2525 |
| *Sup Frontal gyrus, medial* | *R* | *4* | *40* | *54* | *9.07* | |
| Cereb Crus 2 | R | 16 | -78 | -40 | 9.04 | 109 |
| Cereb Lob 6 | R | 18 | -72 | -24 | 7.90 | 123 |
| ITG, posterior | R | 46 | -60 | -12 | 7.76 | 267 |
| Cereb Lob 7b | L | -30 | -70 | -48 | 7.06 | 148 |
| Cereb Lob 8 | R | 34 | -62 | -54 | 6.65 | 102 |
| Cereb Crus 2 | L | -14 | -82 | -40 | 6.51 | 232 |
| *Cereb Crus 1* | *L* | *-20* | *-76* | *-24* | *4.70* | |
| Cereb Lob 6 | L | -34 | -66 | -26 | 6.28 | 96 |
| Cereb Vermis 9 | L | 0 | -60 | -38 | 6.23 | 504 |
| *Cereb Lob 9* | *R* | *12* | *-46* | *-48* | *6.22* | |
| *Cereb Lob 9* | *R* | *14* | *-62* | *-46* | *4.82* | |
| Cereb Crus 1 | L | -40 | -52 | -40 | 6.13 | 20 |
| Cereb Lob 4-5 | L | -6 | -48 | -14 | 5.20 | 52 |
| ITG, posterior | L | -52 | -62 | -10 | 4.51 | 27 |
| IFG triangularis | L | -50 | 36 | 26 | 4.28 | 26 |
| Cereb Crus 1 | L | -46 | -62 | -30 | 3.76 | 13 |

L: left; R: right; MNI: Montreal neurological institute; STG: superior temporal gyrus; IFG: inferior frontal gyrus; MTG: middle temporal gyrus; Supp Motor tracts Area: supplementary motor tracts area; Sup: superior; Cereb Crus: cerebellum crus of ansiform lobule; Cereb Lob: cerebellum lobule; Cereb: cerebellum;; ITG: inferior temporal gyrus.

Supplementary Table 4: Activations, cluster size and coordinates for voice mean energy> normal & F0 flattened voices contrast, whole brain voxelwise *p*<.05 FDR correction, k>10.

| | | MNI coordinates | | | | |
|---|---|---|---|---|---|---|
| Region label | Hemisphere | X | Y | Z | T value | Cluster size (voxels) |
| MTG, posterior | L | -68 | -36 | -6 | 13.54 | 23425 |
| *Supramarginal gyrus* | *R* | *56* | *-2* | *42* | *12.24* | |
| *STS, posterior* | *L* | *-46* | *-42* | *2* | *10.93* | |
| *MTG, posterior* | *R* | *54* | *-34* | *-8* | *10.70* | |
| *Putamen* | *R* | *28* | *-10* | *10* | *10.07* | |
| *STS, posterior* | *R* | *70* | *-34* | *4* | *10.05* | |
| *MTG, posterior* | *R* | *68* | *-38* | *4* | *10.04* | |
| *MTG, posterior* | *R* | *66* | *-42* | *6* | *9.98* | |
| *Rolandic operculum* | *R* | *48* | *-2* | *12* | *9.25* | |
| *Cereb Crus 1* | *R* | *20* | *-70* | *-36* | *8.59* | |
| *Cereb Crus 1* | *L* | *-14* | *-68* | *-30* | *8.37* | |
| *Cereb Crus 1* | *L* | *-22* | *-72* | *-30* | *8.34* | |
| IFG opercularis | L | -60 | 12 | 28 | 7.45 | 516 |
| Precentral gyrus | *L* | *-58* | *2* | *34* | *7.19* | |
| STG, anterior | L | -52 | 4 | -6 | 6.73 | 66 |
| IFG opercularis | R | 50 | 12 | 26 | 6.22 | 65 |
| IFG triangularis | L | -44 | 28 | 10 | 6.06 | 45 |
| Brainstem POTPT | L | -14 | -26 | -14 | 5.15 | 33 |
| STG, mid | R | 52 | -4 | -10 | 4.25 | 34 |
| IFG triangularis | R | 54 | 28 | 26 | 4.16 | 33 |
| MTG, mid | R | 68 | -12 | -20 | 3.85 | 13 |
| Globus pallidus | R | 20 | 2 | -4 | 3.21 | 14 |
| Putamen | R | 22 | 4 | 12 | 2.72 | 10 |

L: left; R: right; MNI: Montreal neurological institute; MTG: middle temporal gyrus; STS: superior temporal sulcus; Cereb Crus: cerebellum crus of ansiform lobule; IFG: inferior frontal gyrus; STG: superior temporal gyrus; Brainstem POTPT: major brainstem motor tracts pathway of the parieto-occipito-temporo-pontine tract.
